# Differential Methylation by Early Life Adversity in the Future of Families Child Wellbeing Study

**DOI:** 10.64898/2026.02.27.708594

**Authors:** Siena Dumas Ang, Samantha Chin, Lisa M Schneper, Rachel A Johnston, Kalsea J Koss, Colter Mitchell, Daniel A Notterman, Barbara E Engelhardt, Catherine Jensen Peña

**Affiliations:** Lewis-Sigler Institute for Integrative Genomics, Princeton University, Princeton, NJ 08544; Department of Computer Science, Princeton University, Princeton, NJ 08544; Department of Molecular Biology, Princeton University, Princeton, NJ 08544; Zoo New England, Boston, MA 02121; Broad Institute of MIT and Harvard, Cambridge, MA 02142; Department of Human Development and Family Science, University of Georgia, GA, 30602; Institute for Social Research, University of Michigan, Ann Arbor, MI, 48104; Gladstone Institutes, San Francisco, CA 94158; Department of Biomedical Data Science, Stanford University, Stanford, CA 94305; Princeton Neuroscience Institute, Princeton University, Princeton, NJ 08544

## Abstract

Early life adversity (ELA) has a well-established link to mental health disorders later in life, yet the molecular mechanisms behind this relationship are incompletely understood. The Future of Families Child Wellbeing Study (FFCWS) provides an opportunity to examine experience encoding in the genome through functional changes in DNA methylation (DNAm) in a cohort enriched in subjects exposed to ELA. We investigated epigenome-wide differences in DNAm across thirteen early-life exposures in salivary samples from FFCWS participants. We calculated differential methylation associations with disease-linked genetic variants, evaluated tissue-specific gene expression, and assessed the persistence of DNAm changes from ages 9 to 15 years. Using data from the mSTARR-seq assay, we characterized methylation-dependent regulatory activity. Differential methylation in the FFCWS validated prior results and identified new genomic regions associated with child adversity. Differential methylation occurs in genomic regions likely to impact gene expression, and affected genes are expressed in disease-relevant tissues. We also identified association of genetic variants associated with downstream disorders near differential methylation, including depression, alcohol and substance use, and anxiety disorders. Overall, cumulative ELA is associated with specific DNAm changes, functional regulation, and persistence over time. Our findings indicate that ELA-associated differential methylation in the FFCWS does not simply occur at random, but in genomic regions that are functional. Our results support the conclusion that altered DNAm represents a biological link between early life experience and later health outcomes.

## Introduction

Early life adversity (ELA) negatively impacts health and behavior across the lifespan^1^. ELA includes a wide range of experiences classically quantified as adverse childhood experiences (ACEs), spanning neglect, abuse, and various aspects of household dysfunction. Exposure to ACEs increases the risk of mental health disorders later in life, including conduct disorder, depression, anxiety, and substance abuse, beginning as early as adolescence^2^. ELA also impacts physical health and increases the risk for cardiovascular disease and metabolic disorders^3^. In turn, altered behavior and health states can undermine physical and socioeconomic outcomes^4–7^. However, the cellular mechanisms through which early experience may simultaneously impart a long-lasting impact on both the body and the mind are incompletely understood.

The epigenome is thought to sit at the interface of developmental biology and environmental experience, such that experiences can become etched into the epigenome to regulate gene expression.^8,9^ While our genetic sequence is mostly fixed, which genes are expressed, as well as the timing and extent of expression, is regulated by DNA methylation (DNAm) and other epigenetic mechanisms. DNAm, the addition of a methyl group to CpG dinucleotides (cytosine followed by guanine), is relatively stable across mitotic cell division and over the lifespan^10,11^. DNAm at transcriptional regulatory regions, including gene promoters, long-range *cis*-regulatory regions, and transcription factor binding sites, serves to regulate gene expression, which in turn influences cellular state and downstream phenotypes^12^.

The methylome has been shown to respond to differences in childhood experience, from socioeconomic status^13,14^ to abuse^15^. The impacts of ELA on DNAm have been measured in large, longitudinal cohorts of children^16–18^, including in the ALSPAC Study^19^, the Dutch Famine Birth Cohort^20^, and the Raine Study^21^. However, these cohorts come from general populations that typically have low occurrences of ELA^16,22^, reducing statistical power to examine the impact of ELA specifically. These studies also include relatively homogeneous, predominately white populations, raising the question of the extent to which models of health risk generalize across ancestral populations^23,24^. This is an important consideration given that many mental and physical health disorders disproportionately affect populations understudied in genomics and health in the United States^25^.

The Future of Families and Child Wellbeing Study (FFCWS) provides an unprecedented opportunity to study differential methylation (DM) and ELA in a large, diverse population. The FFCWS is an expansive, longitudinal dataset following a cohort of children and their families. There has been substantial research in the FFCWS on ELA experiences^26,27^ and adverse outcomes^28^ using the sociological survey data. The FFCWS also includes saliva samples and methylation arrays at ages nine and fifteen. Associations between FFCWS ELA and methylation have been used to validate findings in other studies. For instance, one recent study that focused on ELA and methylation in the ALSPAC study used FFCWS methylation data as a replication cohort.^16^ Another study considered methylation as a mediator between ELA and depressive symptoms in adolescence in the ALSPAC study, and used FFCWS data to replicate these results.^29^ ELA in FFCWS participants has been linked to “methylation age”^30^ in addition to other biomarkers of adversity such as shorter telomere length^31,32^. Other studies have investigated ELA and methylation in the FFCWS, but have focused on single exposures rather than comprehensively defined, longitudinal adverse early-life experiences^33–36^. A major goal of our study was to thoroughly examine the relationship between peripheral DNAm and both cumulative ELA and specific, early-life adverse experiences surveyed in the diverse FFCWS cohort.

While many studies have demonstrated an impact of early-life experiences on DNAm, few have linked those DNAm changes with gene regulation. Empirical studies often find that methylation does not necessarily impact neighboring gene expression^37^. While DM is frequently enriched in areas of the genome harboring genes of interest, most studies fail to investigate disease-appropriate tissue types or validate functional effects experimentally. This is confounded further by the unknown correspondence of samples collected from peripheral tissues, such as saliva, to specific tissues of interest. Thus, there is a need to incorporate experimental regulatory data and cross-tissue expression studies to determine the mechanisms by which DNAm might functionally regulate later life outcomes.

Here, we performed epigenome-wide association studies (EWAS) and identified differentially methylated regions (DMRs) associated with specific measures of ELA in the FFCWS cohort. We measured the relationship between ELA DMRs and genetic disease risk, assessed the impacts of ELA DM in multiple complementary ways, and leveraged the longitudinal aspect of FFCWS to gain insight into the long-term stability of ELA-associated DMRs. Using experimental data from the mSTARR-seq assay, we showed that FFCWS DM is associated with methylation-dependent regulatory activity. Our work elucidates novel insights about how DNAm may functionally mediate the long-term impact of early life adversity.

## Materials & Methods

The FFCWS follows children born in U.S. cities between 1998 and 2000^38^; children born to unmarried mothers were oversampled at a ratio of 3:1, and the cohort contains a large number of Black, LatinX, and low-income families. For the study, families (mother and child, and father when possible) were interviewed at childbirth and again when the children were aged one, three, five, nine, and fifteen years.

### ELA exposures studied in the FFCWS cohort

FFCWS children had exposure to a range of ELA, including adverse family environments and unfavorable health events. To catalogue these experiences, the FFCWS survey asked questions in a variety of categories, including parental involvement, school performance, housing instability, traumatic experiences, and emotional development.^38^ The study also collected salivary DNA samples and processed these samples with genome-wide DNA and methylation arrays.

To study ELA-associated differential methylation (DM), we first generated related exposure variables for FFCWS participants (Table 1, S1), considering only participants that have methylation data and complete survey data at age 9 (see *Supplemental Materials & Methods*). While not all of our variables are typically considered *exposures* (e.g., BMI), we refer to them collectively as “exposures” to maintain consistency. The number of samples included in each analysis and the distribution of scores varied by exposure (Table 1; Figures 1, S1).

**Table 1.**
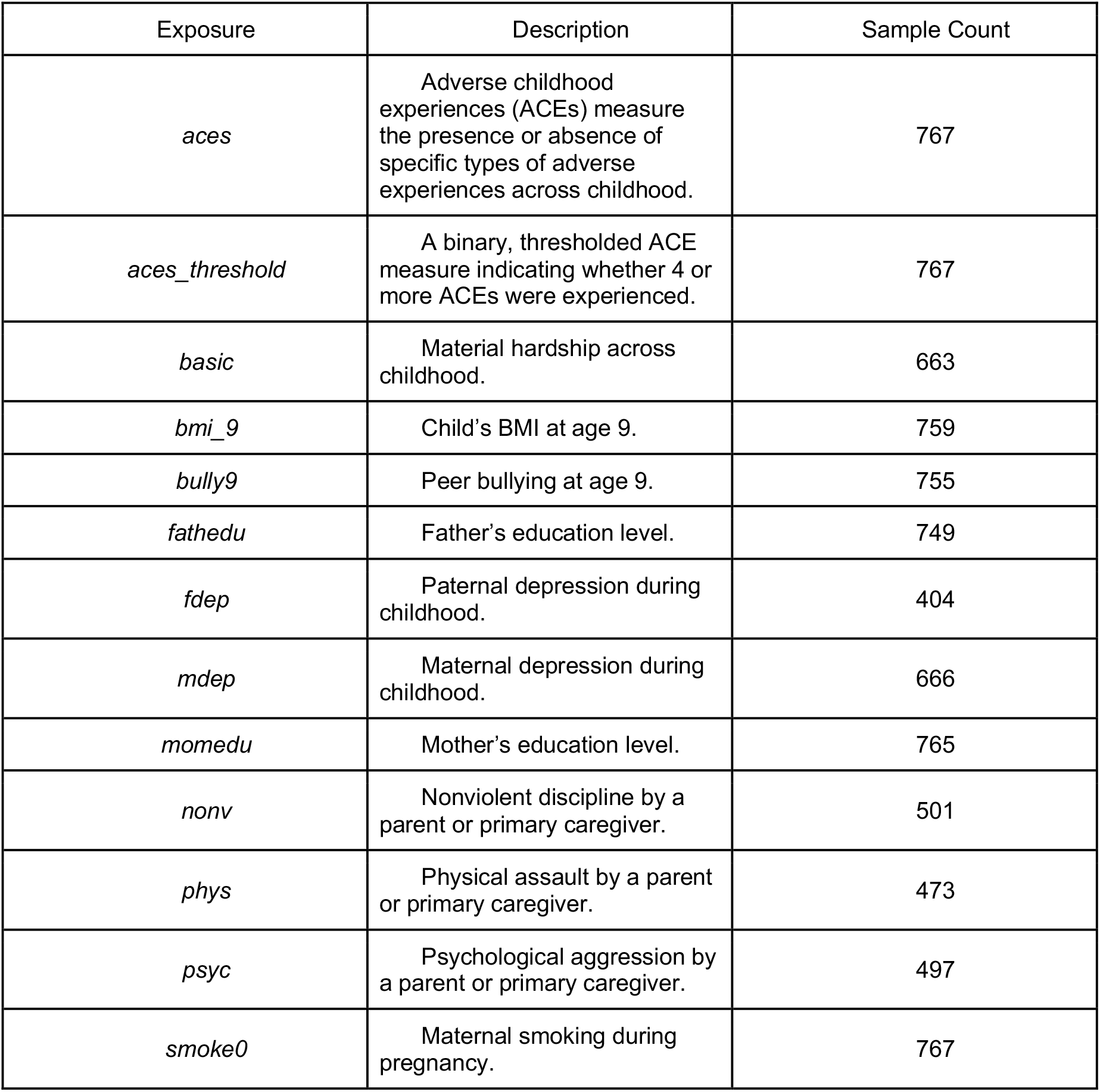
FFCWS ELA variable names (“exposures”), descriptions, and the number of samples with complete exposure, methylation, and genetic data.

| Exposure | Description | Sample Count |
| --- | --- | --- |
| *aces* | Adverse childhood experiences (ACEs) measure the presence or absence of specific types of adverse experiences across childhood. | 767 |
| *aces_threshold* | A binary, thresholded ACE measure indicating whether 4 or more ACEs were experienced. | 767 |
| *basic* | Material hardship across childhood. | 663 |
| *bmi_9* | Child's BMI at age 9. | 759 |
| *bully9* | Peer bullying at age 9. | 755 |
| *fathedu* | Father's education level. | 749 |
| *fdep* | Paternal depression during childhood. | 404 |
| *mdep* | Maternal depression during childhood. | 666 |
| *momedu* | Mother's education level. | 765 |
| *nonv* | Nonviolent discipline by a parent or primary caregiver. | 501 |
| *phys* | Physical assault by a parent or primary caregiver. | 473 |
| *psyc* | Psychological aggression by a parent or primary caregiver. | 497 |
| *smoke0* | Maternal smoking during pregnancy. | 767 |

**Figure 1.**
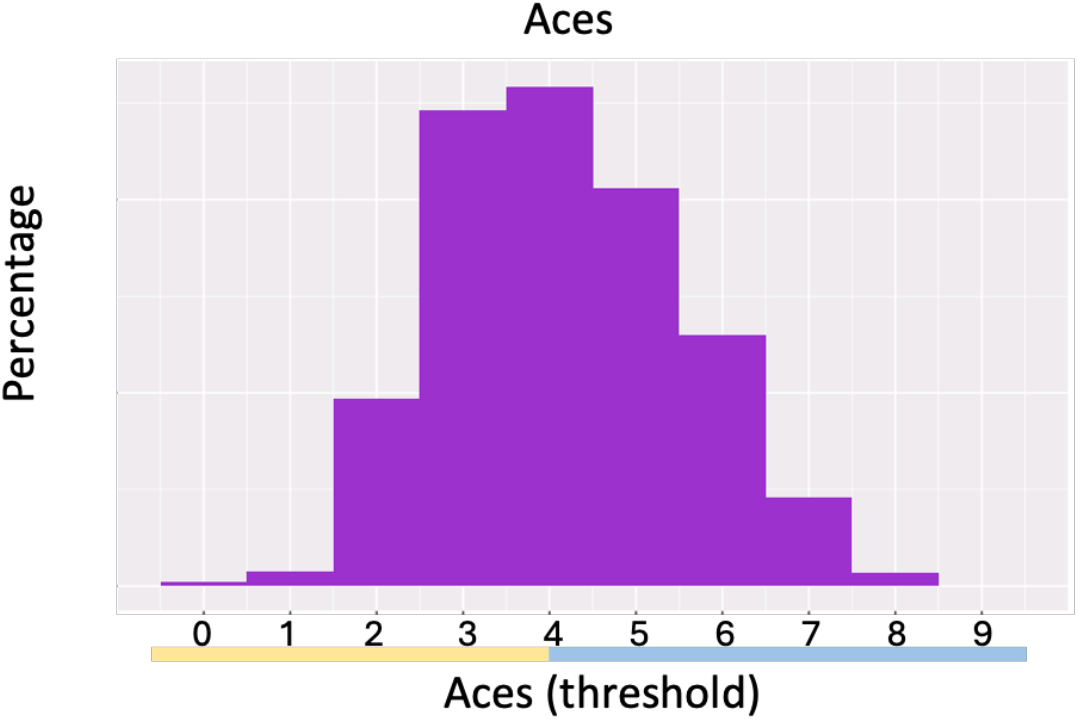
Percentage of cumulative ACE scores (*aces*) across samples. Thresholded cumulative ACEs (*aces_threshold*) are shown in yellow (0) and blue (1) corresponding to whether at least 4 ACEs were experienced. 65% of samples experienced four-or-more ACEs.

Our work considered ELA at a cumulative adverse experience level as well as specific, single-modal adversities across development. Cumulative adverse childhood experiences^28^ (ACEs) reflect the presence or absence of ACE types across childhood, built retrospectively using primary caregiver surveys (*Supplemental Methods*). Experiencing multiple ACEs is associated with negative adult physical and mental health outcomes^2,3,28,39^; these risks are increased with exposure to at least four ACEs^28,40^. As such, we also included a thresholded ACE score by splitting the score into high and low based on whether at least four ACEs were experienced (Figure 1).

### Methylation methods

Saliva samples were collected from FFCWS children at ages 9 and 15 as described in the FFCWS biomarker documentation^41^. Samples were randomized and processed on either the Illumina Infinium Human Methylation450K (450K) or Illumina Infinium MethylationEPIC (EPIC) arrays according to the manufacturer’s protocol. We focus our analyses on the 450K array data, as this earlier array is more consistent with broader literature and existing results^42–44^. As there is minimal overlap (*n=7*) in participants with data from the 450K and EPIC arrays, we include the EPIC array as an independently analyzed cohort (Figures S1-S9).

### Epigenome-wide association study (EWAS)

We identified distinct differentially methylated CpG sites associated with each developmental exposure (Table 1: Exposures), controlling for genetic background, ethnicity, sex, and maternal smoking status during pregnancy with the following equation:

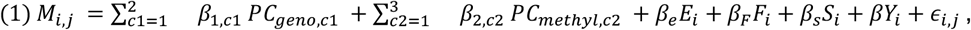

where *M*_*i,j*_ is the M-value (the log ratio of methylated to unmethylated probe intensity signal) of a CpG site for individual *i, i* = 1, . . ., *n* and *j* is the CpG site index; *PC*_*geno*_ is the genetic principal components (PCs) for *c1*=*1-2*; *PC*_*methyl*_ is the methylation PCs for *c2*=*1-3*; *E*_*i*_ is self-reported ethnicity coded as a factor; *F*_*i*_ is a binary variable for sex; *S*_*i*_ is a binary variable representing maternal smoking status during pregnancy and is included in all regressions except when *Y*_*i*_ = *smoke*0; and *Y*_*i*_ is the composite exposure score.

### Identification of differentially methylated regions (DMRs)

Despite including hundreds of participants, statistical power to detect site-level associations is limited by the need to perform multiple testing correction across hundreds of thousands of CpG sites genome-wide. We therefore leveraged the local correlation of methylation across the genome to combine individual CpG sites into differentially methylated regions (DMRs) to improve sensitivity and statistical robustness.^45,46^ Using the *comb-p*^45^ package, we first applied a “moving averages” method of p-value correction, then built DMRs by accounting for spatial correlation and multiple hypothesis testing^47^. While DMRs reflect spatially correlated DM, the magnitude and direction of change may vary at each CpG site within a DMR, precluding direct comparison of uniform increases or decreases in methylation across a region. To generate regions, we required a site q-value of *0*.*05* to begin a region (where a q-value is the Benjamini-Hochberg^48^ corrected p-value, capturing false discovery rate or FDR), and another low q-value site within 1000bp to extend a region. Finally, each region was assigned a Stouffer-Liptak p-value and corrected using Sidak correction^49^, which controls the family-wide error rate (FWER), or the probability of making one or more false discoveries at this corrected p-value threshold. Regions with a Sidak-corrected p-value (*Sidak-corrected p*<*0*.*001)* were deemed DMRs.

### Regression with downstream disorders in genome-wide association studies (GWAS)

We examined overlap of DM with nearby disease-associated single nucleotide polymorphisms (SNPs) using the NHGRI-EBI GWAS Catalog^50^ and dbSNP^51^. Identifiers used in GWAS Catalog queries are included in Data S2. SNPs with any association in the GWAS catalog were filtered to those within a 50/250/500/1000bp window of CpG sites. We calculated associations between FFCWS DM and a binary indicator of whether the nearest SNP was annotated to a disorder known to be affected downstream of ELA (Exposures; Table 1, *Supplemental Methods*) as follows:

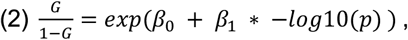

where *G* is a binary indicator for whether the SNP is associated with a disorder of interest and *p* is the uncorrected DM p-value for each CpG site. As these analyses evaluate trends in association strength rather than identifying specific sites or regions, uncorrected p-values were used to preserve the continuous structure of the association signal. We considered an exposure to be associated with a disorder group when *β*_1_ was greater than expected under a one-sided empirical null distribution generated from 15,000 permutations (*p*_*emp*_ < 0.05).

### Methylation-dependent regulatory association

We quantified association of FFCWS DM with methylation-dependent regulatory activity based on published empirical data (measured experimentally) from the mSTARR-seq assay in non-challenged K562 cells^52^. We regressed DM p-values on methylation-dependent function, defined as the log difference of regulatory activity (mSTARR-seq ratios of RNA to DNA) in methylated and sham contexts, using the model:

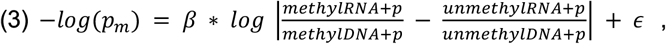

where *p*_*m*_ is the DM p-value for each CpG site, *methylRNA* and *methylRNA* are the mean RNA and DNA counts respectively from mSTARR-seq in the methylated context overlapping the CpG site of interest, *unmethylRNA* and *unmethylRNA* are the mean RNA and DNA counts respectively from mSTARR-seq in the unmethylated context, and *p* is a pseudocount of 20. Empirical p-values were calculated based on a null distribution generated from 20,000 circular-shift permutations of mSTARR-seq regions.

### GTEx tissue expression

Since methylation is highly tissue-specific, we sought to validate our results by determining expression of genes overlapping DMRs in tissues relevant to ELA. Using the Gene-Tissue Expression (GTEx) project^53^, a large database of tissue-specific gene expression, we evaluated the level of expression for genes overlapping DMRs across exposures. We focused on tissues implicated in stress-associated diseases broadly (*Supplemental Methods*). We also tested whether expression of genes overlapping ELA-associated DMRs were expressed greater than or equal to chance using a one-sided Wilcoxon rank-sum test; *q-values* were generated from Benjamini-Hochberg^48^ multiple testing correction.

### Differential methylation at Y15

With the FFCWS variable scores through age 9, we performed an EWAS (Equation 1) with methylation data at age 15. We then computed DMRs with the same method and cutoffs as the age 9 methylation data. We calculated the percentage of DMRs retained at ages 9 and 15, where a DMR is retained if it overlaps with a DMR at the other age.

*Full methods are included in Supplemental Methods*.

## Results

### Differentially methylated regions associated with ELA and other exposures

We identified between 3 and 238 (*Sidak-corrected p<0*.*001*) DMRs associated with each of the FFCWS exposures. We report DMRs and their overlapping genes (Data S1) and documented links to genetically-associated conditions based on prior GWAS results (Table S3) and hand-curated DM literature (Table S4).

To validate our approach, we first examined genes overlapping DMRs linked with BMI at age 9 (*bmi_9*) and maternal smoking during pregnancy (*smoke0*), as these are well-studied variables in prior literature. We mapped genes to GWAS-associated conditions (Table S3) and to broader DM literature (Table S4). Indeed, genes overlapping DMRs for *bmi_9* are associated with body composition traits such as BMI, waist-hip ratio, waist circumference, hip circumference, and body weight. Genes overlapping *smoke0*-associated DMRs have similarly been implicated in GWAS studies of smoking initiation, smoking status, smoking cessation, and nicotine dependence, as well as other outcomes. Further, *MYO1G*^54–59^ and *AHRR*^55–60^ overlap *smoke0*-associated DMRs and have been found to be differentially methylated in children exposed to tobacco prenatally across tissue types throughout the lifecourse—from newborn cord blood to whole blood in midlife (Table S4). Taken together, replication of obesity and maternal smoking-related associations provide evidence that our DMR methods are enriched in biologically relevant genes.

Next, we investigated genes overlapping DMRs associated with other specific ELA exposures. Consistent with the increased risk of developing mental health disorders after ELA, we found FFCWS ELA-associated DMRs in regions of the genome previously linked to psychiatric disorders. In GWAS studies^61^, we found that DMRs associated with ELA exposures in the FFCWS (i.e., *aces, aces_threshold, basic, bully9, fdep, mdep, psyc, phys*) overlapped areas of the genome linked to conditions such as schizophrenia and bipolar disorder, stress response, mood and behavioral disorders, depression, alcohol dependence, or substance abuse (Table S3). DMRs associated with cumulative ACEs (*aces, aces_threshold*) overlapped genes with prior associations to aggressive behaviors, Alzheimer’s disease, asthma and childhood onset asthma, blood pressure measures and hypertension, body composition traits, body height, cognitive function, educational attainment, mood instability, and anxiety-like behavior in animal models^62^. We also identified links with disorders affected downstream of ELA (Table S4), such as aggressive behavior, alcohol use disorder, drug addiction, psychotic experiences in young adulthood, and schizophrenia, providing evidence that behavioral problems may be epigenetically mediated by ELA.

Childhood trauma has been consistently linked to behavioral problems and adult cognitive function^63–68^. In line with this, the gene *TMEM232*, which overlaps both *aces* and *aces_threshold* DMRs, has been linked to mood instability, educational attainment, and cognitive function measurements (Table S3). Notably, ELA-associated DMRs identified in the current analyses contain genes that have been associated with physical abuse and other exposures in the ALSPAC cohort^17,18,69,70^ (Table S4). *NR4A2*, overlapping *nonv-* and *psyc-*associated DMRs, encodes a transcription factor essential for development and function of dopaminergic circuitry of the brain, which has been implicated in psychiatric disease^71–74^ and Parkinson’s^75–77^. The relationship of many well-studied traits in GWAS studies and our DMR results highlight DNAm as a potential molecular mechanism for these relationships. For FFCWS ELA-associated DM not previously connected to adversity, our results provide novel insights implicating ELA. In sum, our findings validate prior work and support the potential role of DNAm as a mediator between ELA and downstream health outcomes.

### ELA-associated differential methylation (DM) is proximal to SNPs implicated in psychiatric disorders

While genes overlapping ELA-associated DMRs have been implicated in psychiatric disease by GWAS and other literature, we sought to quantify this relationship more precisely. We examined the association between exposure-specific DM and nearby disease-associated single-nucleotide polymorphisms (SNPs) using a logistic regression (Equation 2). With annotations from the GWAS Catalog^50^ and SNPs from dbSNP^51^, we focused on SNPs from genome-wide association studies examining mental and substance use disorders that are known risks following ELA. Empirical significance was calculated across 15,000 permutations.

We observed deviation from the empirically-derived null distribution (Figure 2; Data S2; *p*_*emp*_ *< 0*.*05*) of the grouping of all disorders for *bmi_9 (p*_*emp*_*=0*.*002), bully9 (p*_*emp*_*=0*.*001), fathedu (p*_*emp*_*=0*.*004)*, and *psyc (p*_*emp*_*=0*.*013)*. In addition to the grouping of total disorders, *bullying at age 9* (*bully9*) and *psychological aggression* (*psyc*) were associated with the most disorder types: *bully9* with alcohol and substance (*p*_*emp*_=*0*.*049*), anxiety (*p*_*emp*_=*0*.*02*), depression (*p*_*emp*_=0.0001), and *psyc* with alcohol (*p*_*emp*_=*0*.*026*), alcohol and substance (*p*_*emp*_=*0*.*026*), and bipolar disorder and schizophrenia (*p*_*emp*_=*0*.*028*). In addition, *cumulative ACEs* (*aces*) were associated with disordered eating (*p*_*emp*_=*0*.*045*) and *thresholded cumulative ACEs* (*aces_threshold*) with OCD (*p*_*emp*_*=0*.*004*). Additional exposures were linked to anxiety (*momedu, p*_*emp*_*=0*.*023*), bipolar disorder and schizophrenia (*bmi_9, p*_*emp*_*=0*.*018; fathedu, p*_*emp*_*=0*.*002*), depression (*bmi_9, p*_*emp*_*=0*.*002*), OCD (*fdep, p*_*emp*_*=0*.*018*), PTSD (*fathedu, p*_*emp*_*=0*.*028; mdep, p*_*emp*_*=0*.*036*). These findings are consistent with prior evidence linking childhood trauma to depression^28,78–81^, alcohol or substance use^28,40,82,83^, anxiety disorders^81,84,85^, bipolar disorder and schizophrenia^85–90^, disordered eating^82,91,92^, OCD^93,94^, and PTSD^95,96^. At a relaxed empirical significance threshold (*p*_*emp*_ *< 0*.*1*), additional exposures were associated with alcohol (*bully9, p*_*emp*_*=0*.*053*; *momedu, p*_*emp*_*=0*.*09*), alcohol and substance (*momedu, p*_*emp*_*=0*.*096*), anxiety (*mdep, p*_*emp*_*=0*.*052; nonv, p*_*emp*_*=0*.*067*), depression (*phys, p*_*emp*_*=0*.*072*), disordered eating (*momedu, p*_*emp*_*=0*.*059; psyc, p*_*emp*_*=0*.*056*), and PTSD (*nonv, p*_*emp*_*=0*.*07*). *Maternal smoking in utero* (*smoke0*) did not relate strongly to psychiatric disease, with no associations even under a more permissive significance threshold (*p*_*emp*_ *< 0*.*1*).

**Figure 2.**
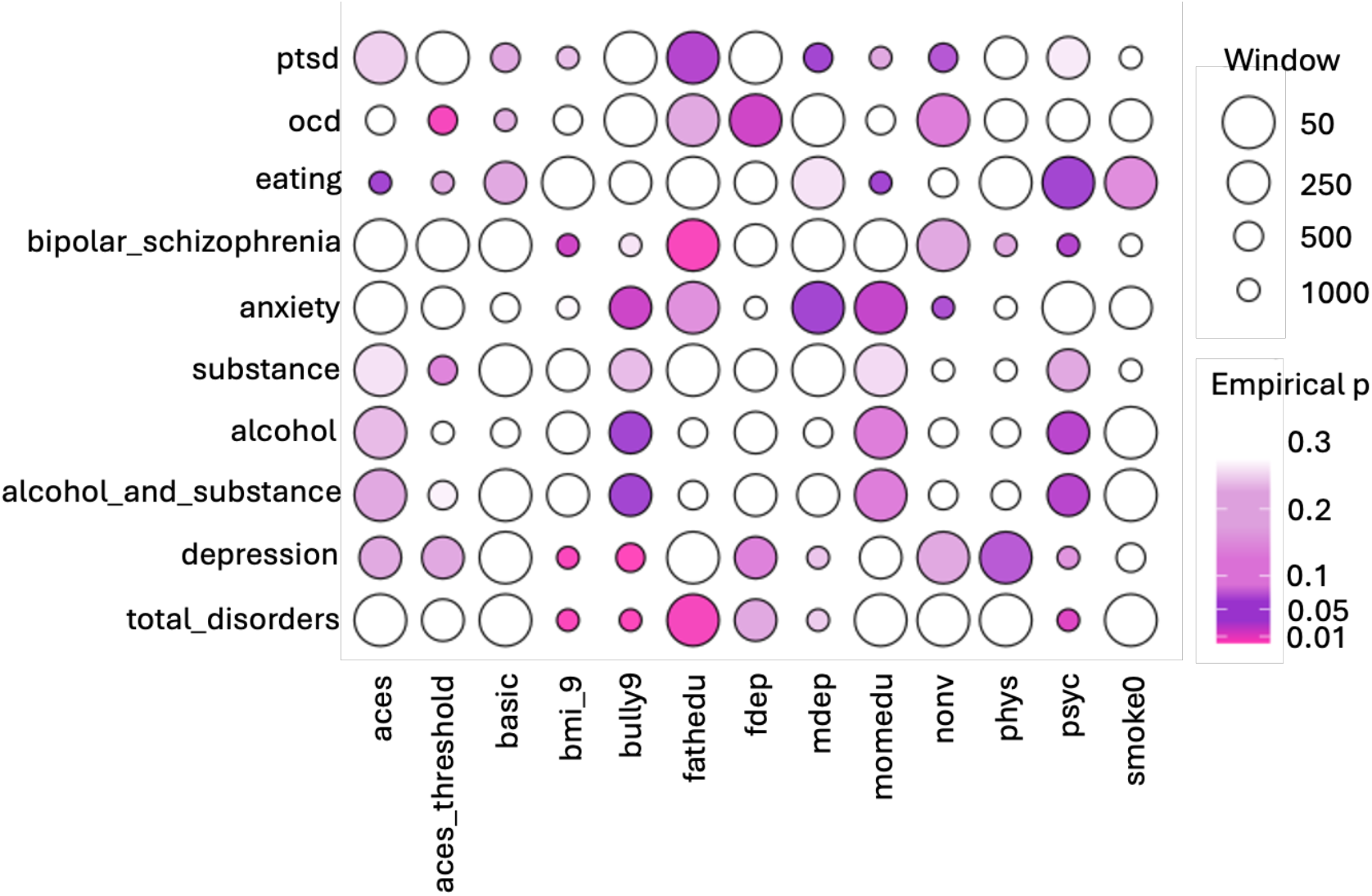
Association of differential methylation with nearby SNPs and relevant disorders. Windows between exposure-associated DM and nearby disease-associated SNPs (within 50/250/500/1000bp) are represented as circle size inversely proportional to window size, such that closer DM-SNP pairs occupy larger circles. Color represents empirical *p-values*: *p*_*emp*_*< 0*.*3* are shown on a light purple gradient, with stronger evidence *p*_*emp*_*< 0*.*05* displayed as darker purple to pink.

One caveat of this approach is the sensitive relationship of SNPs to ancestry; the FFCWS population is considerably more diverse relative to the populations represented in SNP databases^97^. dbSNP integrates data across large-scale sequencing projects like gnomAD^97^; as these resources gain additional ancestral diversity, this analysis should be repeated. Nevertheless, the presence of such associations despite ancestry heterogeneity reinforces a possible role for these sites in biological function and neuropsychiatric disease. Together, across many types of ELA, our results suggest that ELA-associated DM is enriched proximal to loci that have been implicated in psychiatric conditions and adverse downstream health outcomes.

### ELA-associated differential methylation (DM) occurs in areas of the genome where regulatory function is methylation-dependent

While it is tempting to infer the functional impact of DNAm based solely on the location of methylation in the genome, DNAm does not always impact gene expression in empirical studies.^37^ For DM to serve as a mediator of downstream outcomes, it must influence gene regulation. Thus, it is important to distinguish between regions of the genome where methylation affects gene regulation and regions where it does not. However, aside from foundational mSTARR-seq studies^52,98^, methylation-dependent functional analysis has rarely been applied to DM studies.

We sought to quantify methylation-dependent regulatory effects using experimental data from mSTARR-seq^52^, a high-throughput, CpG-free reporter assay^98^ that allows for comparison of regulatory activity of each DNA fragment in the methylated and unmethylated conditions. Significance was calculated by regressing p-values of DM and RNA to DNA ratio comparisons between methylated and unmethylated contexts on a log scale (Equation 3). All exposures except *fdep* and *momedu* showed positive trends with methylation-dependent regulatory activity. Further, *cumulative ACEs, thresholded cumulative ACEs, material hardship, BMI, bullying, father education, maternal depression, physical assault*, and *psychological aggression* were all significantly associated with methylation-dependent regulatory activity (all *p<0*.*05*; Figure 3; full statistics in Table S5). *Bullying, psychological aggression*, and *physical assault* showed the largest magnitude of association with functional DNAm. Our results suggest that ELA-associated DM occurs in areas of the genome where gene expression is methylation-dependent. This is a key advance of our study, demonstrating one mechanism by which ELA may functionally lead to downstream consequences through DNAm.

**Figure 3.**
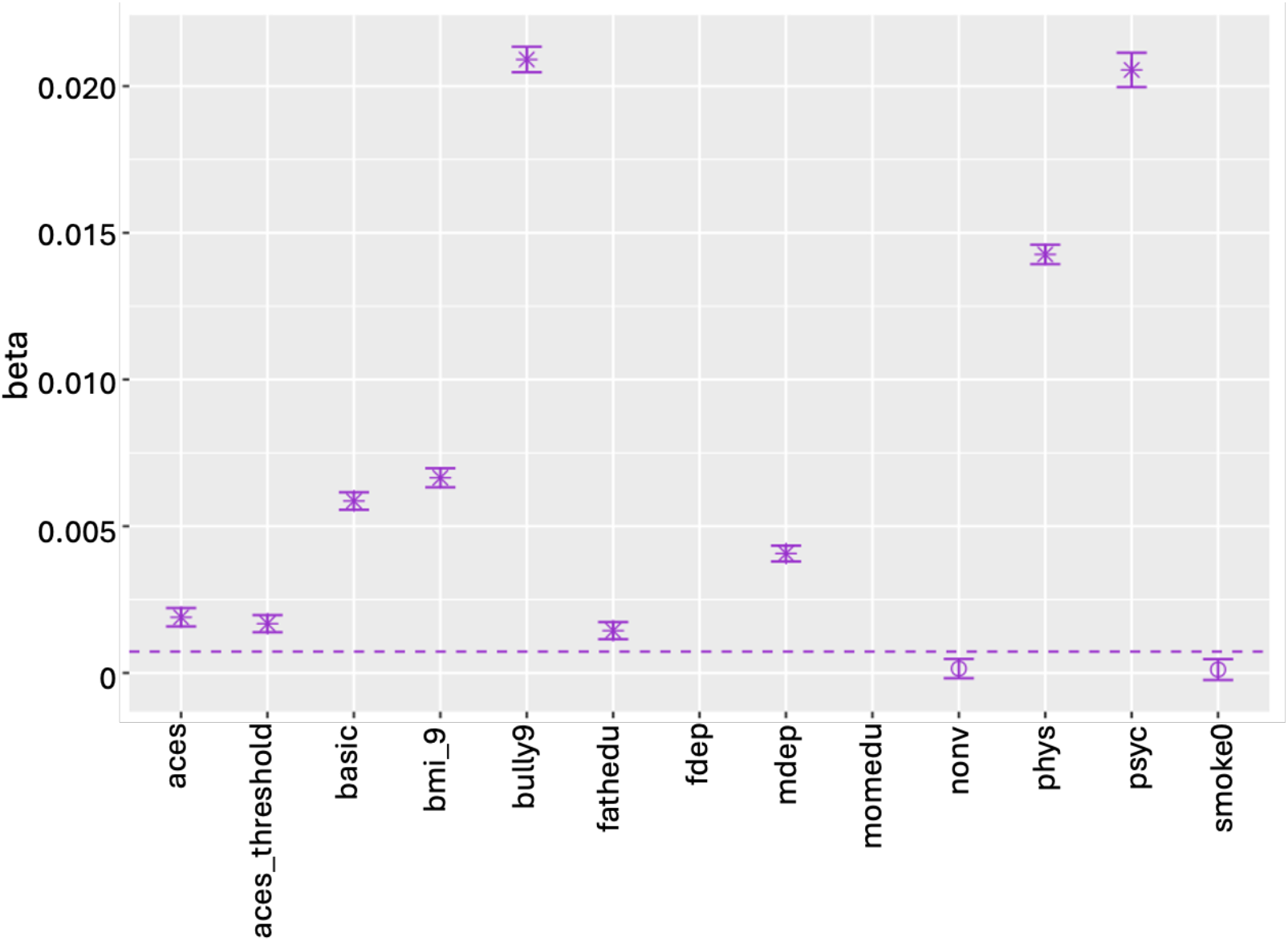
Relationship between differential methylation and methylation-dependent gene regulatory activity. Beta coefficients for each regression between DM significance and methylation-dependent regulatory activity slopes (*β*, Equation 3) for each exposure. All beta slopes above the dashed line met statistical significance (*p*_*emp*_ *< 0*.*05*). Individual statistics are: *aces (β=0*.*0019, p*_*emp*_*=1*.*31e-2), aces_threshold (β=0*.*0017, p*_*emp*_*=1*.*15e-3), basic (β=0*.*0059, p*_*emp*_*=5e-5), bmi_9 (β=0*.*0067, p*_*emp*_*=5e-5), bully9 (β=0*.*0209, p*_*emp*_*=5e-5), fathedu (β=0*.*0014, p*_*emp*_*=4*.*94e-2), fdep (β=-0*.*0152), mdep (β=0*.*0041, p*_*emp*_*=5e-5), momedu (β=-0*.*0065), nonv (β=0*.*0001), phys (β=0*.*0143, p*_*emp*_*=5e-5), psyc (β=0*.*0206, p*_*emp*_*=5e-5), smoke0 (β=0*.*0001)*.

### Tissue-specific expression of DMR-linked genes

FFCWS methylation data was extracted from saliva, a practical and minimally invasive sample to collect from children. Saliva-derived methylation primarily reflects methylation patterns in epithelial and immune cells^99^. However, a concern of this and other EWAS research related to childhood adversity is that both DNAm and gene expression are highly tissue-specific. Saliva-derived DMR-associated genes may or may not even be expressed in tissues of interest. This is a known issue, particularly because EWAS studies often attempt to understand neuropsychiatric and physical health disorders rooted within the brain and other organs. To address this, we mapped tissue-specific expression of genes proximal to ELA-associated FFCWS DMRs using the Gene-Tissue Expression (GTEx) project^53^. Here, we focused on expression within immune, endocrine, reproductive, adipose, cardiovascular, lung, and brain tissues, as these tissues are implicated in adversity-linked diseases^100–107^. We tested expression of gene sets using a one-sided Wilcoxon rank-sum test and stratified expression by transcript levels, where high expression was defined as median transcripts per million (TPM) ≥ 10 (see *Supplemental Methods*).

First, we examined expression of genes overlapping *bmi_9-*associated DMRs (*n=16* genes) across the body, hypothesizing that these genes would be expressed in adipose tissues^108^. Indeed, 62.5% of these genes were expressed in subcutaneous adipose and visceral adipose tissues (Figure 4A) at higher levels than background (*q<0*.*0015* and *q<0*.*0015*, respectively). Genes overlapping *bmi_9-*associated genes were also expressed across multiple brain and reproductive tissues (Data S3), consistent with obesity’s implication in structural brain changes^109^ as well as female infertility^110^. We next examined expression of saliva-derived DNAm among genes overlapping *smoke0-*associated DMRs, expecting expression in lungs^111,112^. Indeed, 82.9% of *smoke0-* associated DMRs (*n=35* genes) were expressed in the lungs at higher than background(*q<2*.*5e-7;* Figure 4B). Gene expression within the frontal cortex of the brain has also been linked with maternal smoking during pregnancy^101^, which is consistent with expression that we found in brain tissues including the frontal cortex (74.3% of genes expressed; *q<3e-8*). (Figure 4B; Data S3). For both *bmi_9-* and *smoke0*-associated DMRs, these findings suggest that genes near saliva-derived DMRs are expressed in tissues validated by prior literature. These findings suggest that tissue expression of genes associated with saliva-derived DMRs from other childhood exposures may also be disease-relevant.

**Figure 4.**
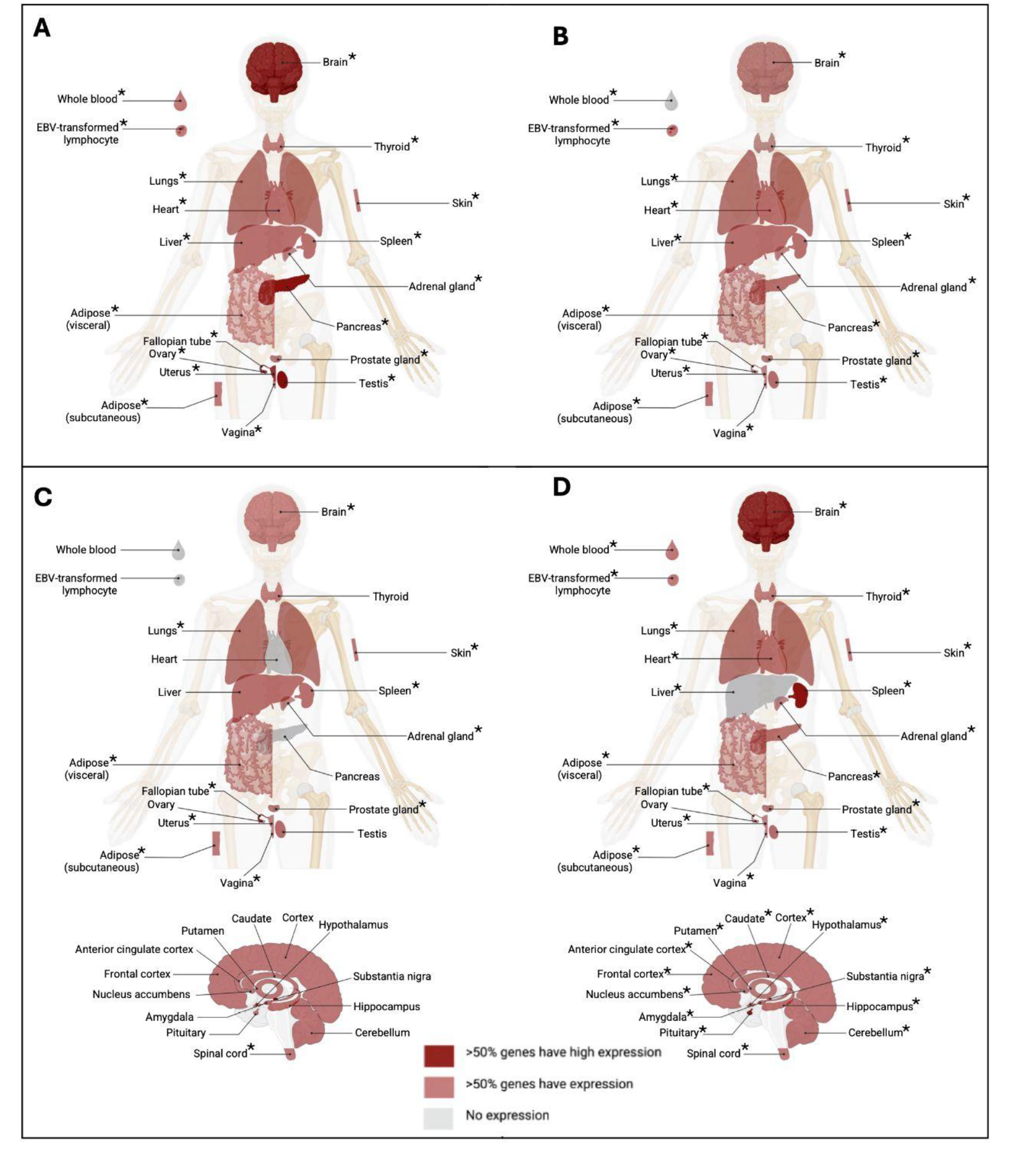
Tissue-specific expression of genes overlapping DMRs. For (A) *bmi_9*: BMI at age 9, (B) *smoke0*: maternal smoking in utero, (C) *aces_threshold*: four or more cumulative ACEs, and (D) *bully9*: peer bullying at age 9, tissues are colored to denote where DMR-associated genes are expressed, based on the GTEx project. Light red indicates that at least 50% of genes are expressed in a given tissue; dark red indicates at least 50% of genes have high expression (median TPM ≥ 10) in a given tissue. * indicates that expression is greater than expected by chance (Wilcoxon rank-sum test) relative to the full gene set included in GTEx (*q < 0*.*05*).

Based on these validations, we sought to characterize where genes overlapping ELA-associated DMRs are expressed. Consistent with prior findings that adverse childhood exposures are associated with a variety of mental^15,113^ and physical health conditions^40,114^, we observed expression across a variety of tissue types (Figure 4C-D; Figure S3). Among the genes overlapping *aces_threshold*-associated DMRs (*n=6* genes), 50% were expressed in the brain (Figure 4C). *SKAP2* and *C8orf31* were expressed in all brain regions; *TMEM232* was expressed in all brain tissues except the cerebellar hemisphere; and *HOXA5* and *HOXA3* were expressed in spinal cord. Genes overlapping *aces_threshold*-associated DMRs were expressed in reproductive tissues, consistent with prior findings that childhood adversity may accelerate reproductive development^115^. Despite the relatively small number of genes, genes overlapping *aces_threshold-*associated DMRs had enriched gene expression in 30 tissues (*q<0*.*05*; Data S3).

Across specific childhood exposures, we found consistent expression of DMR-associated genes in the brain. For example, genes overlapping *bully9*-associated DMRs (*n=5* genes; Data S3) were also expressed throughout the brain and in spleen, lungs, heart, thyroid, adipose, and reproductive tissues (Figure 4D). In addition, DMR-associated genes for all other exposures had expression in brain and other tissues (Figure S3; Data S3). These findings support evidence that methylation derived from saliva samples is potentially relevant to function within brain^116^, endocrine and metabolic physiology^106^, and the maturation and function of the reproductive system^107,115^. Our results suggest the broad applicability of DMRs found in saliva samples by implicating tissues across the body where ELA may be recorded in the epigenome.

### Differential methylation (DM) persists at Y15

The long-term stability of methylation is one way that early life experiences recorded in the epigenome have lasting effects. It is also possible that effects of early life exposures may emerge slowly or prime latent effects, only appearing as new DMRs in later years. To evaluate the long-term effects of childhood exposures, we calculated DMRs from salivary methylation collected at age 15 and assessed whether DMRs were present at both ages, only at age 9, or only at age 15. For consistency, we used the same ELA exposure scores through age 9 and performed the same EWAS and region generation with methylation data at age 15 (Equation 1), replacing *M*_*i,j*_ with M-values at age 15 (Data S4).

Broadly, we observed that, on average (median), 33.3% of exposure-specific DMRs at age 9 were maintained at age 15, and 36.8% of DMRs at age 15 were also present at age 9 (Figure 5; Table S6). DMRs associated with thresholded cumulative ACEs were maintained at the highest level, and no new *aces_threshold*-associated DMRs were found at age 15—indicating persistent effects of DM in response to thresholded cumulative ELA.

**Figure 5.**
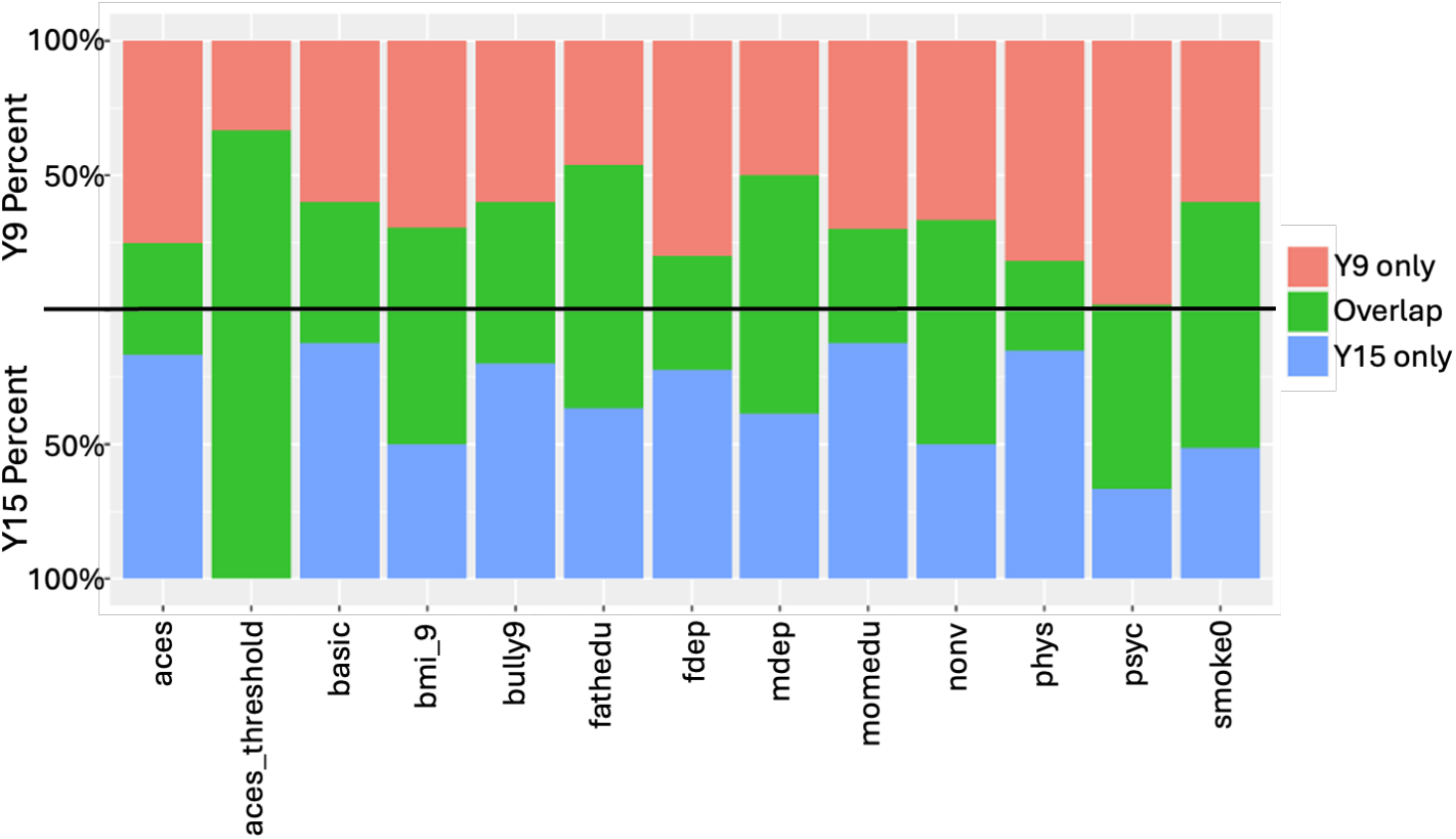
Percent of DMRs lost, retained, or added between ages 9 and 15. Values on top reflect percent of Y9 DMRs specific to Y9 (red) or overlapping with Y15 (green). Values on bottom reflect percent of Y15 DMRs specific to Y15 (blue) or overlapping with Y9 (green). Each exposure is represented as a separate bar.

While DNAm can be highly stable, it is not expected that all DMRs overlap completely. DNAm analysis from the ALSPAC cohort identified time-dependent changes in CpG methylation that were considered either stable, emergent, latent, or primed, and only about 50% of DM that was concordant across a similar time frame from ages 7 to 15^16^. Similarly, we observed some loss of year 9 DMRs associated with each exposure (Figure 5). Loss of early-life DMRs associated with ELA could be attributed to simple fading with time or to normal age-related methylation changes finally “catching up” by adolescence. There is growing evidence for ELA to accelerate epigenetic development^117–123^. Additional longitudinal studies incorporating more densely sampled measurements across development are necessary to distinguish this possibility in the FFCWS cohort.

We also identified DMRs at age 15 that were not present at age 9 (*aces, basic, bully9, fathedu, fdep, mdep, momedu*, and *phys*; Figure 5; Table S6). This may be due to several possibilities, including: delayed impact of the exposure, effects of exposures that emerged slowly across development, accumulation of additional exposures since age 9, or latent epigenetic priming^18,57,124^. Indeed, earlier work identified DNAm associated with ELA that did not emerge until 15 years of age^16^. Previous studies have also found evidence that ELA can prime stress sensitivity^125–127^ or alter the HPA-axis^128–130^ leading to downstream molecular changes that may only emerge with additional stress exposures. Together, these results suggest that while many methylation changes are stable across development, particularly for children exposed to four-or-more ACEs, DMRs may also be relatively dynamic, impacting different genomic regions at different times.

### Replication on the EPIC array

We next investigated DM associated with ELA in the FFCWS for a cohort (*n=1093*) processed on the EPIC array. As there were only seven samples shared between the two arrays, the EPIC array was treated as an out-of-sample cohort. While exposure scores were created with the same method, the methylation data was processed independently, leading to distinct DMRs from the EPIC data. All analyses can be found in the Supplement (Table S6, Figures S4-S7). There is prior evidence of weak correlation between the 450K and EPIC arrays at individual CpG sites in samples collected from children^131^; in addition, cell-type composition in saliva collected from children has interindividual heterogeneity^132^. We found few explicitly overlapping DMRs for the same exposure on both arrays (1 overlapping for *bmi_9* and 11 for *smoke0*). However, DMRs are shared across exposures (Figure S2). While we observed general concordance in broad patterns across analyses of GTEx tissue expression (Figures S3-4) and persistence of DMRs (Figures 5, S7), we did not observe similar concordance in tests for proximity of nearby SNPs (Figures 2, S5) or methylation-dependent regulatory associations (Figures 3, S6). This may also be related to differences in data distribution across samples included in each array, in particular differences in the self-identified ethnicity of samples (Figure S8), which may contribute to lack of methylation consistency^13,14,133,134^. Continued sampling in future studies across diverse populations will help resolve these differences.

## Discussion

This study is the first to examine specific ELA-associated DM epigenome-wide in the FFCWS, which has both a large sample size and a high incidence of ELA, with a specific eye toward understanding the potential downstream impact of DNA methylation. Our DMRs validate prior literature and identify new genomic regions associated with different aspects of childhood adversity. Several genes overlap DMRs associated with multiple exposures and disorders, including *TMEM232* (associated with *aces* and *aces_threshold*) and *NR4A2* (associated with *nonv* and *psyc*). DMRs were implicated in relevant conditions such as depression, consistent with substantial prior literature showing increased risk of psychiatric conditions following ELA^28,29,102,125,135–137^. Our findings support the growing literature suggesting that DNAm represents a biological link between early life experiences and later life health outcomes.

Here, we applied genomic analyses to investigate the potential impact of ELA-associated DNAm, including functional relevance. Our findings demonstrate that ELA-associated DM in the FFCWS occurs near genetic variants associated with downstream disorders identified through GWAS. Drawing on experimental data, we find that childhood exposures are associated with DNAm in areas of the genome where alterations are likely to impact gene expression. This distinctive aspect of our study addresses a major gap in the literature as to whether DM, particularly in intergenic regions of the genome, is likely to have any impact on gene expression. Our findings support the potential relevance of DM reported in prior EWAS studies and set the stage for causal investigation in future studies. We also identified tissues where DMR-associated genes are expressed, validating the predictive relevance of salivary DNAm sampling for disease-related tissue types across the body^138,139^. This is important given the caveat that epigenome regulation is often tissue specific, but access to disease-relevant tissues from living human subjects, especially children, is limited. Finally, we leveraged the longitudinal nature of FFCWS to investigate the long-term effects of ELA experiences on DNAm, as past studies have shown that some DNAm is persistent throughout the lifecourse^59,70,140,141^, while others show loss of early-life DM^142^ or delayed emergence of DM^143–145^. While approximately one third of DMRs were stable from childhood to adolescence, late emergence of new DMRs at age 15 reflects the dynamic nature of the epigenome and has been suggested as a contributing factor in why complex disease may unfold over many years^16^.

Although there is generally a lack of reproducibility across studies of cumulative ELA^146,147^, our study identified ELA-associated DMRs near genes that were also identified in the ALSPAC study, including validating a link with physical abuse^69^ and other ACE-related exposures^17,18,69,70^ (Table S4). While the majority of DMR-associated genes were distinct from recent findings, lack of replication across cohorts may reflect differences in population ancestry between the FFCWS and more homogeneous white populations examined in prior studies^16,69^. Recent work that used FFCWS as a replication cohort was only able to partially replicate loci association direction with childhood adversity discovered in the ALSPAC study^16^. Similarly, in our study, there were differences in the self-reported ethnicity of the FFCWS samples processed on the 450K versus EPIC arrays, in addition to array-related differences in chemistry and probe coverage. Even within our own FFCSW cohort, we find that these factors likely lead to critical differences in the regions identified.

A surprising finding in our results was the large variation in the number of DMRs across exposures. In particular, *psychological aggression* had an unexpectedly large number of DMRs. This could be related to normalization of the FFCWS methylation data, as FFCWS DNAm was preprocessed prior to this manuscript for consistency across studies and includes probes overlapping SNPs as well as low variance sites that may affect quantification of methylation levels. The *psyc* score represents averages across questions and timepoints, resulting in a near-continuous variable whose distribution appears symmetric relative to other similarly constructed exposures (Figure S1). One possibility is that residual, probe-level structure might interact with patterns in exposure distribution, which could lead to weakly significant, spatially correlated DM sites that are aggregated into DMRs. Nevertheless, many genes overlapping *psyc-*associated DMRs have been implicated by GWAS and other literature in depression, bipolar disorder and schizophrenia, and other psychiatric, behavioral, and physical health disorders, suggesting some veracity of these results.

Another important consideration for the current study is that the cumulative ACE scores (*aces, aces_threshold*) used in this study were extrapolated from questionnaires administered to children and parents, rather than the original ACE questionnaire, which is typically asked retrospectively in adulthood. Many of the responses came from parents rather than children themselves, and may therefore be underreported. These differences should be noted when comparing to studies that explicitly use the ACE questionnaire asked in adulthood. However, all traditional exposures were included in order to build our cumulative ACE scores (*aces, aces_threshold*), making the FFCWS cumulative ACEs a good proxy for the standard ACE score.

In sum, these findings demonstrate the potential for ELA-associated DM to shape later life health outcomes. Across adverse childhood exposures, our results support the interpretation that ELA-associated DM in the FFCWS is specific to regions that are functionally relevant in disease-related tissues. This research lays the groundwork for future studies to more thoroughly investigate associations with specific ELA exposures in other diverse cohorts, build sophisticated models of joint contributions to methylation between specific forms of ELA, quantify similarities in epigenetic signatures across ELA, and causally investigate methylation-dependent regulatory activity. Through a thorough characterization of ELA-associated DNA methylation in rich study data, our work adds a unique perspective on the embedding of ELA exposures in the epigenome and how these mechanisms may influence later life outcomes.

## Acknowledgements

This work was funded by the National Science Foundation Graduate Research Fellowship Program (SDA); Graduate Fellowships for STEM Diversity, formerly National Physical Sciences Consortium sponsored by the National Security Agency (SDA); New York Stem Cell Foundation (CJP); R01-HD076592 (DAN); R01-HD103669 (DAN); R01-MH103761 (CM); and R01-MD011716 (CM). CJP is a New York Stem Cell Foundation Robertson Investigator and is also funded by NIH R01MH129643. BEE is a CIFAR Fellow in the Multiscale Human Program and is funded by the Parker Institute for Cancer Immunology (PICI), the Chan-Zuckerberg Institute (CZI), the Biswas Family Foundation, NIH NHGRI R01 HG012967, and NIH NHGRI R01 HG013736. Sara McLanahan founded the FFCWS and passed before publication; we remember her brilliant mind, passion for research, and generous spirit in this work. We thank Jenny Tung for input on study design and analyses. Thank you to Sarah Kocher for discussion and advice; William-Sebastian Gil for previous version gene lookups; Bob Dumas for reading early versions of the manuscript; and Hiei Rose, along with Mykonos and Maui Dumuchi Rosang, for their support. Parts of some figures were generated with BioRender.

## Author contributions

SDA and BEE designed the study with additional input from CJP, DAN, and RAJ. SDA performed the analyses along with SC. DAN, LMS, RAJ, KJK, and CM contributed data and analysis pipelines: DAN and LMS generated and processed the methylation and genetic data; KJK made the cumulative ACE score; and CM created all other composite scores. The manuscript was written by SDA, SC, and CJP, with input from all authors.

## Data availability

DMRs for the 450K and EPIC arrays are available in Data S1.

