## Supplemental Methods, Tables, Figures for "Differential Methylation by Early Life Adversity in the Future of Families Child Wellbeing Study"

*Pages*

**Supplemental Methods** 2-6

**Supplemental Tables:** 7-23
Table S1: FFCWS ELA variable names, descriptions, and counts included in the EPIC array
Table S2: Identifiers used in GWAS Catalog queries
Table S3: Annotations of genes to GWAS conditions for genes that overlap with FFCWS ELA-associated DMRs
Table S4: Annotations of genes overlapping FFCWS ELA-associated DMRs to DM genes in prior literature
Table S5: Statistics of methylation-dependent regulatory associations by exposure
Table S6: Percent of DMRs lost, retained, and arising between Y9 and Y15

**Supplemental Figures:** 24-31
Figure S1: Exposure distribution scores for the 450K and EPIC array
Figure S2: Percent of DMRs that overlap with another DMR across the 450K and EPIC array
Figure S3: Tissue-specific expression of genes overlapping 450K array DMRs for additional exposures
Figure S4: Tissue-specific expression of genes overlapping EPIC array DMRs for additional exposures
Figure S5: Enrichment of differential methylation with nearby SNPs and associated disorders (EPIC array)
Figure S6: Relationship between differential methylation and methylation-dependent gene regulatory activity for 450K and EPIC arrays
Figure S7: Percent of DMRs lost, retained, or added for the EPIC array between ages 9 and 15
Figure S8: Percent of self-described ethnicity across participants on 450K and EPIC array

**Supplemental Data (indexed only):**
Data S1: DMRs for 450K and EPIC arrays
Data S2: Regression test statistics for regression with GWAS Catalog (*empirical p-values*) and GWAS identifiers for disorder groupings
Data S3: Percentages of tissue-specific gene expression across exposures and Wilcoxon rank-sum test statistics for tissue-specific expression (*q-values*)
Data S4: Y15 DMRs for 450K and EPIC arrays

**References** 32-35

### Supplemental Methods

*Study sample characteristics*

FFCWS families are comprised of 4,898 children born in major cities in the United States between 1998 and 2000.[1] The FFCWS cohort mothers were recruited in the hospital during childbirth. Baseline interviews were conducted with mothers in the hospital soon after the child's birth, and fathers, when available, were interviewed in the hospital or by phone. Follow-up interviews were conducted when the child was one, three, five, nine, and fifteen years old.

Cumulative ACEs measure how many different ACE types were experienced from ages 1 through 9, recording whether each type of ACE was experienced across physical abuse, emotional abuse, sexual abuse, physical and emotional neglect, maternal interparental violence, parental mental illness, parental substance use, parental incarceration, and parental divorce. Information on these variables was taken from mother, father, primary caregiver, and child surveys. Individual early life composite variables were created from survey questions measured from ages 1 to 9 for the following phenotypes: parenting scores--*physical assault* (*phys*), *psychological aggression* (*psyc*), *nonviolent discipline* (*nonv*); parental depression--*maternal depression* (*mdep*), *paternal depression* (*fdep*); and *material hardship* (*basic*). The following variables document life events recorded at baseline: parental education--*mother education* (*momedu*), *father education* (*fathedu*); and *maternal smoking during pregnancy* (*smoke0*). *Bullying in year 9* (*bully9*) is an aggregated score based on the child's self-reporting of adverse peer experiences at age 9. We also include the adjusted z-score of the child's body mass index at year 9 (bmi_9) as a well-studied phenotype to serve as a benchmark for our methods. Details on how each individual score was constructed are below.

*ELA composite scores*

Each individual was given a composite score for thirteen phenotypes based on a set of relevant FFCWS survey questions. We require responses at all applicable timepoints for a given sample and phenotype.

*Parenting Scores*

Parenting phenotypes were aggregated over responses at child ages 3, 5, and 9. The question scales come from the Parent Child Conflict Tactics Scales, and asked the primary caregiver how frequently they had done certain actions. Participants self-scored either a 1 (once), 2 (twice), 3 (3-5 times), 4 (6-10 times), 5 (11-20 times), or 6 (more than 20 times in the last 12 months).

*Physical assault* (*phys*) measures how frequently a child was spanked, hit on the bottom with a hard object, slapped, pinched, or shaken by a parent or primary caregiver. Physical assault ranges from 0-6 based on frequency of physical assault by parent or primary caregiver across ages 3, 5, and 9 (Figure S1B).

*Psychological aggression* (*psyc*) is scored based on how often a parent yelled at their child, threatened to spank the child, swore at them, called the child dumb or lazy, or threatened to send away or kick the child out of the house. Psychological aggression ranges from 0-6 and measures average over childhood of psychological aggression and verbal abuse (Figure S1C).

*Nonviolent discipline* (*nonv*) is measured by how frequently a parent explained why something was wrong, gave the child something else to do instead of what he/she was doing, took away privileges from or grounded the child, or put their child in a “time out”. Scores range from 0-6, after averaging across ages 3, 5, and 9 parental self-scoring of nonviolent discipline parenting frequency (Figure S1A).

*Parental Depression*
Parental depression measures from the mother and father evaluated parental depression when their children were ages 1, 3, 5, and 9. The Composite International Diagnostic Interview (Short Form) was used to determine whether parents met the liberal criteria for depression. Samples were given a score of 0 or 1 at each timepoint and depression scores were averaged across timepoints.

*Maternal depression* (*mdep*) is an average of maternal depression during childhood. Maternal depression ranges from 0-1, and was averaged across ages 1, 3, 5, and 9 (Figure S1E).

*Paternal depression* (*fdep*) is an average of paternal depression during childhood. Paternal depression ranges from 0-1, and was averaged across childhood based on depression criteria at each age (Figure S1D).

*Parental Education*
Parental education was recorded at baseline. Participants self-reported education and received a score of 1 (less than high school), 2 (high school degree/GED), 3 (some college), or 4 (college degree).

*Mother education* (*momedu*) is the mother's education level. Mother education ranges from 1-4 (less than high school education to college degree) recorded at birth (Figure S1H).

*Father education* (*fathedu*) is the father's education level. Father education ranges from 1-4 (less than high school education to college degree) recorded at baseline (Figure S1G).

*Material Hardship*
Material hardship was aggregated over responses at child ages 1, 3, 5, and 9. For each question, participants received a binary score of 0 (No) or 1 (Yes).

*Material hardship* (*basic*) is averaged over whether, in the past 12 months, a child's family received free food, did not pay the full rent/mortgage, got evicted, did not pay full gas/oil/electric bill, or if anyone in the house did not get needed medical attention because of cost. Material hardship ranges from 0-1, and is averaged across age and hardship type (Figure S1F).

*Bullying*
Based on the child's self-reporting of multiple questions at age 9, bullying responses were recorded as 0 (not once in the past month), 1 (1-2 times in the past month), 2 (once a week), 3 (several times per week), or 4 (every day).

*Bullying at age 9* (*bully9*) is based on how often kids in their school or neighborhood picked on or said mean things to them, hit them, took their things without asking, or purposely left them out of activities. Bullying ranges from 0-4 based on the average incidence of bullying experienced at 9 years old (Figure S1I).

*BMI*
BMI measures the child's constructed body mass index Z-score.

*BMI at year 9* (*bmi_9*) is the adjusted z-score of the child's BMI. BMI measures the adjusted z-score of child body mass index (bmi_9) measured at age 9 (Figure S1J). The 450K array included 759 samples and the EPIC array included 1085 samples.

*Maternal Smoking*
Based on self-reporting at baseline, mothers were asked whether they smoked during pregnancy and received a score of 1 (any smoking during pregnancy) or 0 (no reported smoking during pregnancy.)

*Maternal smoking during pregnancy* (*smoke0*) is a binary score of maternal smoking in the prenatal period. Maternal smoking scores range from 0-1 based on whether any smoking in the prenatal period was reported at birth (Figure S1K).

*ACEs*
Adverse childhood experiences (ACEs) measure the presence or absence of a variety of experiences across childhood. In the ACE survey[2,3], the ACE categories span abuse, neglect, and household challenges. The FFCWS adverse childhood experiences (*aces*) score ranges from 0-9, measuring how many adverse experiences were present during childhood (Figure S1L). Each ACE that was coded as “Yes” is counted as one. The aces variable is the cumulative sum across all ACEs: physical abuse, emotional abuse, sexual abuse, physical and emotional neglect, maternal interparental violence, parental mental illness, parental substance use, parental incarceration, and parental divorce.
Physical abuse was coded as “Yes” if the child was spanked by the mother in the past month at age 1; any items on the abbreviated physical assault subscale of the parent-child Conflict Tactics Scale (CTS) were endorsed in the past year at ages 3, 5, or 9 ; or CPS was contacted due to primary caregiver reporting physical abuse at any age, recorded at ages 5 and 9.
Emotional abuse was recorded as “Yes” if any items on the abbreviated psychological aggression subscale of the CTS were endorsed in the past year at ages 3, 5, or 9.
Sexual abuse was coded as “Yes” if the primary caregiver reported sexual abuse as a reason for CPS contact at any age, recorded at ages 5 and 9.
Physical and emotional neglect was recorded as “Yes” if the child met the criteria for physical neglect, emotional neglect, or neglect. Since the CPS questions reported for neglect but did not differentiate between emotional and physical neglect in the FFCWS surveys, physical and emotional neglect are coded as a single ACE. Neglect is recorded if there was an endorsement of child hunger at ages 1, 3, or 5; presence of neglect on the CTS items at ages 3, 5, or 9; or endorsement of CPS questions at ages 5 or 9.
Maternal interparental violence was coded as “Yes” if mothers reported any physical violence from a current new partner, or from the child’s biological father at present or in the last month of an ended relationship. Physical violence is recorded if the child’s biological father or current partner slaps or kicks mother, or hits her with fist or object, at ages 1, 3, 5, or 9; if the partner threw something at them or pushes, grabs or shoves them; if the mother reported she was cut, bruised or seriously hurt in a fight with the child’s father or current partner at ages 5 or 9; or if the mother experienced unwanted sexual acts at ages 1,3, 5, or 9.
Parental mental illness is recorded as “Yes” if the mother or father had a case of major depression at ages 1, 3, 5, or 9, or generalized anxiety disorder at ages 1 or 3.
Parental substance use was coded as “Yes” if either the mother or father had any illegal drug use (including marijuana and hard drugs) or problematic drinking (5+ drinks in a day at age 1 or 4+ at ages 3, 5, and 9).
Parental incarceration was coded as “Yes” if either mothers or fathers were reported as incarcerated at ages 1, 3, 5, or 9.
Parental divorce was recorded as “Yes” if mothers reported a divorce from the child’s biological father at any of the assessments at ages 1, 3, 5, and 9.

*ACEs Threshold*
We also consider a binary, thresholded ACE score. *Aces_threshold* is 0 when three or fewer ACEs are experienced, and 1 when there are four or more ACE exposures.

*Methylation methods*

Samples were collected using Oragene DNA Self-Collection Kits, and DNA was extracted, purified, and stored at -80C. Genomic DNA was genotyped using Illumina PsychChip arrays versions 1.0 and 1.1 and processed according to De Vito et al[4]. The EZ-96 DNA Methylation Kit was used for bisulfite conversion from genomic DNA. Genotype and methylation arrays were performed for 851 children on the 450K, and 1260 children on the EPIC array at age 9.

*M-value generation*

Methylation levels were quantified for each CpG site as M-values, the logarithm of the ratio between the methylated and unmethylated signal.[5] QC was performed with EWAStools[6] to remove probes with detection values (detp) above 0.01 for 450K and 0.05 for EPIC. The ENmix[7] command QCinfo was then used with a 4 bead cutoff and default parameters to remove outlier samples. Additional normalization was done with Enmix using the preprocessENmix command followed by rcp.

*Epigenome-wide association study*

Using methods similar to De Vito et al.[4], we associated relevant CpG site M-values with our composite phenotypes of interest, adjusting for population structure, batch effects, sex, self-reported ethnicity, and maternal smoking during pregnancy. As a specific goal of the FFCWS was to survey a diverse population, we adjusted for population structure by performing principal components analysis (PCA) on sample genetic data.[8] We explored whether the first two PCs related to self-reported ethnicity and included these PCs in our regression to capture the samples' genetic ancestry. We also ran a PCA on the methylation data and included the first three PCs as covariates to account for batch effects[9].

We included self-reported ethnicity as a covariate in our model in order to control for known cultural differences in Black vs. Latinx vs. White households and communities, such as attitudes toward mental health, substance abuse, and physical and verbal punishment[10]. For our adversity-related phenotypes, we controlled for maternal smoking during pregnancy due to evidence that DNA is differentially methylated in children after in utero exposure[11]. As is common practice in genome-wide association studies, we fit the following linear regression model for our continuous, ordinal, and binary phenotypes.

We fit the regression using the R package *limma*[12] from Bioconductor[13] and shrank the standard errors using the empirical Bayes function. This approach generated p-values quantifying differential methylation at each CpG site for each of our phenotypes. Similarly, we performed epigenome-wide association studies using the data from Illumina's EPIC chip to provide an out-of-sample dataset.

*GWAS Catalog associations for genes overlapping ELA-associated DMRs*

We queried the GWAS Catalog with unique trait identifiers (Table S1) using the 9/30/2025 GWAS catalog release made with the *gwascat*[14] package.

*Regression with downstream disorders in GWAS Studies*

We filtered all SNPs in dbSNP to those that had any association in the GWAS Catalog version 2025-09-04. Considering all CpG sites proximal to those SNPs, we assigned a score of 1 to the CpG site if a proximal SNP (within 50/250/500/1000bp) is associated with a disorder of interest, otherwise the site receives a score of 0. SNP locations were based on the hg19 dbSNP locations in the 2025-01-15 version, downloaded here: https://ftp.ncbi.nlm.nih.gov/snp/latest_release/. Associations were based on GWAS Catalog version 2025-09-04.

We performed a logistic regression with an independent variable of the negative log(pvalue) for each site, and whether the nearest SNP was associated with our disorders of interest as the dependent variable. Empirical *p-values* were generated based on a null distribution of 15,000 permutations. To preserve spatial correlation of nearby CpG sites, permutation tests were performed using a block-based randomization of site DM p-values. For each chromosome, probes were ordered by genomic position, circularly phase-shifted, and partitioned into blocks of 200 probes. Blocks were randomly permuted to assign p-values to each CpG site and test statistics were generated by regressing the permuted DM p-values with the disorder binary indicators.

*Methylation-dependent regulatory association*

We used previously published mSTARR-seq data derived from non-challenged K562 cells[15]. The experimental data include RNA and DNA counts across replicates in both methylated (6 replicates) and unmethylated (5 replicates) contexts. We used UCSC's LiftOver[16] to liftover the hg38 regions of the genome to hg19.

For each 600bp region of the genome, we calculated the mean of the DNA replicates after removing the highest and lowest outliers; regions where less than 2 replicates had nonzero DNA counts were omitted. We calculated the mean of the RNA replicates after removing the highest and lowest outliers. There were no requirements on non-zero replicate counts. We filtered regions that had a greater than 25% difference between pseudocount-adjusted DNA means in methylated and unmethylated contexts.

We calculated a regression *p-value* for each exposure associated with methylation-dependent regulatory activity. Empirical p-values were calculated by comparing the observed *p-value* to a null distribution generated from 20,000 circular-shift permutations. mSTARR-seq data were circularly shifted by a random offset within each chromosome in order to preserve the spatial correlation of regions.

*GTEx tissue expression*

We visualized GTEx gene expression levels across different body systems for the set of genes associated with each exposure. Brain tissues have been associated with poverty, maternal smoking during pregnancy, and psychological abuse.[17–19] Immune tissues have been linked to multiple forms of adversity, such as bullying, childhood family instability, and economic insecurity.[20,21] Past studies have established a relationship between the lungs and both in-utero nicotine exposure[22] and material hardship[23]. Childhood adversity also has impacts on neural, endocrine, immune, and metabolic physiology[24], in addition to negatively affecting the maturation and function of the female reproductive system[25].

We labeled a gene as expressed in a tissue if it has at least 0.5 median transcripts per million (TPM), and we classified that expression as high if median TPM ≥ 10. We omitted pseudogenes and genes with mean expression < 0.5 TPM across all GTEx tissues. We included immune tissues (spleen, skin, EBV-transformed lymphocytes, whole blood), endocrine and reproductive tissues (thyroid, pancreas, adrenal gland, liver, pituitary, fallopian tube, ovary, uterus, vagina, testis, prostate gland), brain tissues (amygdala, anterior cingulate cortex, caudate, cerebellum, cortex, frontal cortex, hippocampus, hypothalamus, nucleus accumbens, putamen, spinal cord, substantia nigra), adipose tissues (visceral and subcutaneous), heart, and lungs. For each tissue type, we determined whether at least 50% of the genes overlapping significant DMRs met the baseline expression threshold or the high expression threshold. If at least 50% of the genes for the phenotype met the expression criteria, the tissue was colored light red; if at least 50% of genes met the high expression criteria, the tissue was colored dark red. Tissues were left gray for those where fewer than 50% of genes were expressed. For tissues with more than one sub-tissue type (e.g., skin contains both sun-exposed and non-sun-exposed; heart contains both atrial appendage and left ventricle), the tissue was colored based on the maximum of the sub-tissue types.

#### Supplemental Tables

Table S1. FFCWS ELA variable names, descriptions, and counts included in the EPIC array. Variable names ("exposures"), descriptions, and number of samples with complete exposure, methylation, and genetic data on the EPIC array.

| Exposure | Description | EPIC Sample Count |
|---|---|---|
| aces | Adverse childhood experiences (ACEs) measure the presence or absence of specific types of adverse experiences across childhood. | 1093 |
| aces_threshold | A binary, thresholded ACE measure indicating whether 4 or more ACEs were experienced. | 1093 |
| basic | Material hardship across childhood. | 922 |
| bmi_9 | Child's BMI at age 9. | 1085 |
| bully9 | Peer bullying at age 9. | 1074 |
| fathedu | Father's education level. | 1051 |
| fdep | Paternal depression during childhood. | 567 |
| mdep | Maternal depression during childhood. | 923 |
| momedu | Mother's education level. | 1091 |
| nonv | Nonviolent discipline by a parent or primary caregiver. | 666 |
| phys | Physical assault by a parent or primary caregiver. | 627 |
| psyc | Psychological aggression by a parent or primary caregiver. | 666 |
| smoke0 | Maternal smoking during pregnancy. | 1093 |

Table S2. Identifiers used in GWAS Catalog queries.

| GWAS Condition | Experimental Factor Ontology<br>MONDO Disease Ontology<br>Ontology of Biological Attributes<br>Human Phenotype Ontology |
|---|---|
| ADHD | adhd (EFO_0003888)<br>cognitive_inhibition_measurement (EFO_0007969) |
| Age at Menarche | age_at_menarche (EFO_0004703) |
| Aggressive Behaviors | aggressive_behavior (EFO_0003015)<br>aggressive_behavior_quality (OBA_1000376) |
| Alcohol Use<br>Alcohol Consumption<br>Alcohol Dependence | alcohol_dependence (MONDO_0007079)<br>alcohol_drinking (EFO_0004329)<br>alcohol_depedence_measurement (EFO_0007835)<br>alcohol_use_disorder_measurement (EFO_0009458)<br>alcohol_consumption_quality (OBA_1000840) |
| Alzheimer's Disease<br>Late-onset Alzheimer's Disease<br>Alzheimer's Biomarkers | alzheimers_disease (MONDO_0004975)<br>late_onset_alzheimers (EFO_1001870)<br>alzheimers_biomarker_measurement (EFO_0006514)<br>early_onset_alzheimers (EFO_0022957) |
| Antisocial Behaviors<br>Social Inhibition/Deprivation | social_inhibition_quality (OBA_1000278)<br>borderline_personality_disorder (HP_0012076)<br>social_deprivation (EFO_0009696)<br>social_communication_impairment (EFO_0005427) |
| Anxiety<br>Anxiety Measures<br>Anxiety Disorder<br>Panic Disorder<br>Worry Measurement | anxiety (HP_0000739)<br>anxiety_disorder (EFO_0006788)<br>anxiety_disorder_measurement (EFO_0007795)<br>anxiety_measurement (EFO_0009863)<br>generalized_anxiety_disorder (EFO_1001892)<br>panic_disorder (EFO_0004262)<br>worry_measurement (EFO_0009589) |
| Asthma<br>Childhood Onset Asthma | asthma (MONDO_0004979)<br>childhood_onset_asthma (MONDO_0005405) |
| Autism Spectrum Disorder | autism_spectrum_disorder (EFO_0003756) |
| Autoimmune Disease | autoimmune_disease (EFO_0005140) |
| Behavioral Disinhibition<br>Behavior (Atypical) | atypical_behavior (HP_0000708)<br>behavioral_disinhibition_measurement (EFO_0006946) |
| Bipolar Disorder | bipolar_disorder (MONDO_0004985)<br>bipolar_I_disorder(EFO_0009963) |
| Birth Weight<br>Birth Length<br>Spontaneous Preterm Birth | body_height_at_birth (EFO_0006784)<br>birth_weight (EFO_0004344)<br>spontaneous_preterm_birth (EFO_0006917) |

| | |
|---|---|
| Blood Pressure<br>Systolic Blood Pressure<br>Diastolic Blood Pressure<br>Hypertension | systolic_blood_pressure (EFO_0006335)<br>diastolic_blood_pressure (EFO_0006336)<br>hypertension (EFO_0000537) |
| Body Composition<br>BMI<br>Body Fat Percentage<br>Waist-Hip Ratio (BMI-adjusted)<br>Body Weight | bmi (EFO_0004340)<br>body_composition (EFO_0005106)<br>body_fat_percentage (EFO_0007800)<br>body_fat_distribution (EFO_0004341)<br>waist_hip_ratio (EFO_0004343)<br>bmi_adj_waist_hip_ratio (EFO_0007788)<br>body_weight (EFO_0004338) |
| Body Height | body_height (OBA_VT0001253) |
| Brain Measurements<br>Brain Volume | brain_volume (OBA_2050009)<br>brain_physiology_trait (OBA_VT0015058) |
| Cognitive Function<br>Intelligence | cognitive_function (EFO_0008354)<br>intelligence (EFO_0004337) |
| Coronary Artery Disease | coronary_artery_disease (EFO_0001645) |
| Depressive Symptoms<br>Mood Disorders | mood_disorder (EFO_0004247)<br>depressive_symptom_measurement (EFO_0007006) |
| Diabetes<br>Type 2 Diabetes<br>Insulin Measurements | type_2_diabetes (MONDO_0005148) |
| Drug Use<br>Drug Dependence<br>Heroin Dependence<br>Opioid Dependence | drug_use_measurement (EFO_0007010)<br>heroin_dependence (EFO_0004240)<br>opioid_dependence (EFO_0005611)<br>opioid_use_measurement (EFO_0009937) |
| Early Life Stress<br>Childhood Trauma | childhood_trauma_measurement (EFO_0007979) |
| Eating Behavior | binge_eating (EFO_0005924)<br>eating_behavior (EFO_0007829) |
| Educational Attainment<br>Self-Reported Educational Attainment | educational_attainment (EFO_0011015)<br>self_reported_educational_attainment (EFO_0004784) |
| Emotional Symptoms<br>Mood Instability | mood_disorder (EFO_0004247)<br>emotional_symptom_measurement (EFO_0007803)<br>mood_instability_measurement (EFO_0008475) |
| Insomnia | insomnia (EFO_0004698)<br>insomnia_measurement (EFO_0007876) |
| Major Depressive Disorder | major_depressive_disorder (MONDO_0002009) |
| Neuroticism | neuroticism_measurement (EFO_0007660)<br>neurotic_disorder (EFO_0004257) |

| | |
|---|---|
| Obsessive Compulsive Disorder | ocd (EFO_0004242)<br>obsessive_compulsive_symptom_measurement (EFO_0007802) |
| Personality Disorder | borderline_personality_disorder (HP_0012076) |
| Psychotic Symptoms | psychotic_symptoms (EFO_0005940) |
| Schizophrenia | schizophrenia (MONDO_0005090) |
| Smoking<br>Smoking Status<br>Smoking Initiation<br>Nicotine Use<br>Nicotine Dependence | smoking_behavior (EFO_0004318)<br>smoking_initiation (EFO_0005670)<br>smoking_status_measurement (EFO_0006527) |
| Stress-related Disorders<br>Stress Reaction<br>Response to Trauma<br>PTSD | stress_related_disorder (EFO_0010098)<br>ptsd_symptom_measurement (EFO_0008535)<br>ptsd (EFO_0001358) |
| Substance Use<br>Substance Abuse<br>Substance Dependence | substance_use (MONDO_0002491) |
| Suicide Behavior | attempted_suicide (EFO_0004321)<br>suicidal_ideation (EFO_0004320)<br>suicide_behavior_measurement (EFO_0006882) |
| Testosterone | testosterone_measurement (EFO_0004908) |
| Wellbeing Measurement | wellbeing_measurement (EFO_0007869) |

Table S3. Annotations of genes to GWAS conditions of interest for genes that overlap with FFCWS ELA-associated DMRs across exposures. Unless otherwise cited, all annotations were queried from the NHGRI-EBI GWAS Catalog[27].

| GWAS Conditions | DMR Genes by FFCWS Exposure |
|---|---|
| ADHD | bmi_9 (*MFSD6L*)<br>bully9 (*TYW3*)<br>fdep (*ZFP57*)<br>mdep (*CD40*)<br>psyc (*ANKK1, NTM, WSCD2, IGF1R, RAI1, SREBF1, SPPL2C, MAPT-AS1, ASPSCR1, NGEF, CACNA2D2, IQCJ-SCHIP1, ZFP57, ELFN1, SDK1, PTPRN2, CHD7*) |
| Age at Menarche | psyc (BET1L, WSCD2, GALNT9, IGF1R, SLC38A10, GREB1, CAMK4, PTPRN2) |
| Aggressive Behaviors | aces_threshold (SKAP2)<br>bmi_9 (ABAT)<br>psyc (NTM, OPCML, INHBB, LINC01101, SDK1) |
| Alcohol Use<br>Alcohol Consumption<br>Alcohol Dependence | basic (LRMDA)<br>fdep (APOB)<br>mdep (PRDM8)<br>momedu (NDUFAF6)<br>nonv (NR4A2, ZNF577)<br>phys (HOXC4, PSMD9)<br>psyc (CASZ1, MUC2, OPCML, NR4A2, KCNJ11, HOXC4, PRDM8, NOS3, ANKK1, PRDM16, CRTAC1, OSBPL5, NTM, WSCD2, GALNT9, ZNF280D, MAPT-AS1, ASPSCR1, SNRPD3, SPON2, RGS12, SDK1)<br>smoke0 (NDUFAF6, PRDM16, CHAD, ACSF2, ZBTB38, CMYA5) |
| Alzheimer's Disease<br>Late-onset Alzheimer's Disease<br>Alzheimer’s Biomarkers | aces (TMEM232)<br>aces_threshold (TMEM232)<br>bmi_9 (IKZF4, ABAT, TNXB, FOXP4, PLEC)<br>bully9 (TYW3)<br>fathedu (TNXB)<br>fdep (KLF6, APOB)<br>momedu (ESPNP, LIAT1, NDUFAF6)<br>phys (AKR7A3, OPLAH)<br>psyc (DRAXIN, HSPG2, TGFB2, MGMT, INPP5A, PTDSS2, CHID1, MUC6, OSBPL5, CCND1, NTM, OPCML, SNHG14, CACNA1H, TCF25, SREBF1, SPPL2C, MAPT-AS1, ERCC2, ANKRD30BP2, CECR7, PDE6B, IDUA, DGKQ, SLC26A1, NELFA, PDCD6, CAMK4, TNXB, SDK1, |

| | |
|---|---|
| | BMPER, HECW1, TRIM56, PRKAR2B, EPHA1-AS1, PTPRN2, PLEC, JPH3, CERS3, GSE1)<br>smoke0 (SHANK2, ANK1, NDUFAF6) |
| Antisocial Behaviors<br>Social Inhibition/Deprivation | bully9 (TYW3)<br>fdep (APOB)<br>mdep (CD40)<br>psyc (NTM, WSCD2, IGF1R, ASPSCR1, IQCJ-SCHIP1, ELFN1, SDK1, PTPRN2, CHD7, CASZ1, KCNJ11) |
| Anxiety<br>Anxiety Measures<br>Anxiety Disorder<br>Panic Disorder<br>Worry Measurement | basic (LRMDA)<br>bmi_9 (TNXB)<br>fathedu (TNXB, HLA-DQB2)<br>fdep (APOB)<br>mdep (PRDM8)<br>psyc (DIP2C, WSCD2, PRDM8, PTPRN2, ANKK1, SDK1, OPCML, FOXK2, PENK, NOS1, MAPT-AS1, TCEA2, MGMT, SNHG14, NTM, RGS12, TNXB)<br>smoke0 (SHANK2, CHAD, ACSF2) |
| Asthma<br>Childhood Onset Asthma | aces (BOLL, TMEM232)<br>aces_threshold (TMEM232)<br>bmi_9 (IKZF4)<br>fathedu (SBNO2, HLA-DQB2)<br>fdep (TRPC3, BRD2)<br>momedu (HLA-DPA1, HLA-DPB1)<br>nonv (ZNF577)<br>psyc (PRDM16, GATA3, SH3PXD2A, MUC2, LRP1, SNHG14, GREB1, TUBGCP6, TRPC3, CAMK4, ARHGEF37, GRM4, SDK1, BMPER, CYP1B1-AS1, CYP1B1, SCHIP1, IQCJ-SCHIP1)<br>smoke0 (PRDM16, ZBTB38, CMYA5) |
| Autism Spectrum Disorder | bmi_9 (TNXB)<br>fathedu (TNXB)<br>fdep (ZFP57, BRD2)<br>psyc (DRAXIN, SPPL2C, MAPT-AS1, NGEF, CACNA2D2, ZFP57, EHMT2, TNXB) |
| Autoimmune Disease | momedu (HLA-DPA1)<br>psyc (GATA3, ADCY7, CAMK4, EHMT2) |
| Behavioral Disinhibition<br>Behavior (Atypical) | fdep (ZFP57)<br>psyc (OSTM1-AS1, OSTM1, ZFP57) |
| Bipolar Disorder | bmi_9 (GNA12, PLEC)<br>fathedu (HLA-DQB2)<br>fdep (APOB)<br>mdep (CD40)<br>nonv (ZNF577)<br>psyc (PC, WSCD2, RPS27P25, JPH3, PIPOX, NGEF, |

| | |
|---|---|
| | TUBGCP6, CACNA2D2, CD47, PDE5A, PLEC)<br>smoke0 (SHANK2, PIPOX, RFTN2, MYO1G) |
| Birth Weight<br>Birth Length<br>Spontaneous Preterm Birth | bmi_9 (AMZ1, GNA12)<br>fathedu (ZNF512B)<br>fdep (ZFP57)<br>psyc (CCND1, IGF1R, ADCY7, RAI1, ASPSCR1, LINC01101, PDE5A, ZFP57, NOS3, GPR20, SDK1)<br>smoke0 (ZBTB38, ANK1) |
| Blood Pressure<br>Systolic Blood Pressure<br>Diastolic Blood Pressure<br>Hypertension | aces_threshold (SKAP2, HOXA-AS3, HOXA3)<br>basic (LRMDA)<br>bmi_9 (UNC45A, TNXB, FOXP4, AMZ1, GNA12, PLEC, BRSK2, IKZF4)<br>bully9 (SPTBN4)<br>fathedu (ZNF512B, TNXB, NBR2, HLA-DQB2)<br>fdep (APOB, HOXA-AS2, HOXA-AS3, HOXA3, ZFP57)<br>mdep (UNC45A, RGL3, PRDM8, CD40)<br>momedu (HLA-DPA1, NDUFAF6)<br>phys (NEAT1, HOXC4, PSMD9, OPLAH)<br>psyc (PRKCZ, PRDM16, CASZ1, TGFB2, CRTAC1, PKD2L1, SH3PXD2A, INPP5A, CHID1, KCNJ11, CCND1, NTM, OPCML, SSPN, HOXC4, LRP1, RPS27P25, MSI1, BCL7A, HHIPL1, CACNA1H, UBE2I, GSE1, MAPT-AS1, MAP2K2, TCF7L1, LINC01101, TCEA2, CACNA2D2, RGS12, PRDM8, PDE5A, AHRR, EHMT2, TNXB, GRM4, PACSIN1, ELFN1, SDK1, TRIM56, NOS3, AGAP3, SDCBP, GPR20, PLEC, SMURF2, SCAP, CAMK4, ZFP57, AGER, COL11A2, TAPBP, PTPRN2, GALNT9)<br>smoke0 (PRDM16, ZBTB38, ENPEP, AHRR, ANK1, NDUFAF6, GTF2H4) |
| Body Composition<br>BMI<br>Body Fat Percentage<br>Waist-Hip Ratio (BMI-adjusted)<br>Body Weight | aces (CCDC169-SOHLH2, SOHLH2, BOLL)<br>aces_threshold (SKAP2, HOXA-AS3, HOXA3)<br>basic (LRMDA)<br>bmi_9 (BRSK2, IKZF4, CRYGEP, GNA12, AMZ1, MPRIP, TNXB, PLEC)<br>fathedu (SBNO2, TNXB, HLA-DQB2)<br>fdep (IFI44L, APOB, BRD2, KLF6, ZFP57, HOXA-AS3, HOXA3)<br>mdep (PRDM8)<br>nonv (ZNF577)<br>phys (HOXC4)<br>psyc (CASZ1, DIP2C, GATA3, CRTAC1, MGMT, INPP5A, CHID1, KCNJ11, CCND1, NTM, OPCML, LRP1, WSCD2, RPS27P25, BCL7A, GALNT9, ANKLE2, OTUD7A, ZNF280D, IGF1R, CERS3, CACNA1H, GSE1, JPH3, KLHDC4, GLP2R, RAI1, SREBF1, ANKRD13B, AOC2, ASPSCR1, SLC16A3, CSNK1G2, MAP2K2, GALNT5, |

| | |
|---|---|
| | TCEA2, CBS, TTLL8, SCAP, CACNA2D2, RGS12, PRDM8, PDE5A, AGER, TAPBP, GRM4, PACSIN1, ELFN1, SDK1, ZNF316, NOS3, AGAP3, PTPRN2, CHD7, SSPN, CSHL1, HOXC4, DGKQ, TGFB2, AHRR, PRDM16, MAPT-AS1, FOXK2, PC, MSI1, UBE2I, ELMO3, SPPL2C, GH1, LINC01101, SLC26A1, ZFP57, EHMT2, TNXB, PLEC, PDCD6, SLC12A7)<br>smoke0 (MEX3A, LTBP3, SHANK2, GABRB3, KLHDC4, RFTN2, ZBTB38, ENPEP, CMYA5, NLGN2, AHRR, PRDM16, POU5F1) |
| Body Height | aces (CCDC169, CCDC169-SOHLH2)<br>aces_threshold (SKAP2, HOXA5, HOXA-AS3, HOXA3)<br>basic (PMF1, PMF1-BGLAP, LRMDA, BTBD17)<br>bmi_9 (ACY3, IKZF4, ABAT, MPRIP, TNXB, FOXP4, AMZ1, GNA12, PLEC)<br>bully9 (TYW3)<br>fathedu (SBNO2, TNXB, HLA-DQB2)<br>fdep (IFI44L, KLF6, APOB, TRPC3, ZFP57, BRD2, HOXA-AS3, HOXA3)<br>mdep (TNNC2, CD40, CSNK1E)<br>momedu (ESPNP, HLA-DPA1, HLA-DPB1, NDUFAF6)<br>nonv (LGALS3BP, NR4A2)<br>phys (PSMD9, BLCAP)<br>psyc (PRKCZ, PRDM16, MEGF6, CASZ1, HSPG2, PODN, TGFB2, DIP2C, GATA3, GATA3-AS1, CRTAC1, PKD2L1, SH3PXD2A, MGMT, INPP5A, BET1L, NLRP6, PTDSS2, CHID1, CDKN1C, OSBPL5, MRGPRG-AS1, PC, C11orf86, CCND1, ANKK1, NTM, OPCML, LINC00942, SSPN, LRP1, WSCD2, RPS27P25, MSI1, BCL7A, GALNT9, ANKLE2, HHIPL1, TECPR2, OTUD7A, ZNF280D, IGF1R, CERS3, LMF1, CACNA1H, IFT140, TMEM204, ADCY7, GSE1, JPH3, GLP2R, RAI1, SREBF1, PIPOX, GIT1, ANKRD13B, MAPT-AS1, CSHL1, GH1, SMURF2, TMEM235, LGALS3BP, SLC38A10, ASPSCR1, FOXK2, CSNK1G2, MAP2K2, ISYNA1, GREB1, CYP1B1-AS1, CYP1B1, TCF7L1, INHBB, LINC01101, NR4A2, GALNT5, SLC40A1, ALPI, NGEF, NFS1, TCEA2, CBS, GGT1, SUN2, SCAP, CACNA2D2, IDUA, SPON2, MIR943, NELFA, RGS12, PDE5A, TRPC3, PDCD6, AHRR, SLC6A3, ARHGEF37, UNC5A, ZFP57, HLA-V, EHMT2, TNXB, RNF5, COL11A2, TAPBP, GRM4, PACSIN1, TTYH3, SDK1, ZNF316, BMPER, HECW1, PRKAR2B, NOS3, AGAP3, SMARCD3, CHPF2, PTPRN2, PENK, CHD7, GPR20, RHPN1, PLEC)<br>smoke0 (PRDM16, MEX3A, LTBP3, SHANK2, GABRB3, NLGN2, PIPOX, CHAD, ACSF2, ZBTB38, ENPEP, AHRR, DMGDH, GTF2H4, POU5F1, PIWIL2, ANK1, NDUFAF6) |

| | |
|---|---|
| Brain Measurements<br>Brain Volume | bmi_9 (AMZ1, GNA12, PLEC)<br>mdep (PM20D1)<br>psyc (PRKCZ, PRDM16, SH3PXD2A, LINC00942, MSI1, MAPT-AS1, CSHL1, GH1, IQCJ-SCHIP1, PLEC, HSPG2, PIPOX, GIT1, AGAP3)<br>smoke0 (PRDM16, PIPOX, SEZ6) |
| Cognitive Function<br>Intelligence | aces (TMEM232)<br>aces_threshold (TMEM232)<br>basic (LRMDA)<br>bmi_9 (IKZF4, TNXB)<br>bully9 (ZSCAN12P1)<br>fathedu (TNXB)<br>fdep (ZFP57)<br>mdep (PM20D1, CD40)<br>phys (PSMD9)<br>psyc (MUC2, OPCML, WSCD2, NOS1, LMF1, CACNA1H, RAI1, MAPT-AS1, SCAP, CACNA2D2, IQCJ-SCHIP1, RBM46, SLC12A7, ZFP57, TNXB, SDK1, PTPRN2, CHD7, DIP2C, MGMT, NTM, RPS27P25, GIT1, SPPL2C, FEM1AP3, PRKAR2B)<br>smoke0 (RFTN2) |
| Coronary Artery Disease | bmi_9 (TNXB, ZNF727)<br>fathedu (TNXB)<br>fdep (KLF6, APOB)<br>mdep (RGL3, PRDM8)<br>momedu (NDUFAF6)<br>psyc (PRDM16, CRTAC1, PKD2L1, BET1L, OPCML, LRP1, HHIPL1, IFT140, TMEM204, RAI1, SREBF1, ANKRD13B, MAPT-AS1, CSNK1G2, CYP1B1-AS1, CYP1B1, NFS1, SCAP, RGS12, JAKMIP1, PRDM8, TNXB, ELFN1, TRIM56, NOS3)<br>smoke0 (PRDM16, NDUFAF6) |
| Depressive Symptoms<br>Mood Disorders<br>(see below for MDD) | bmi_9 (TNXB)<br>bully9 (TYW3)<br>fathedu (TNXB)<br>fdep (APOB)<br>Mdep (CSNK1E24)<br>phys (BLCAP)<br>psyc (SSPN, MAPT-AS1, DIP2C, ANKK1, NOS1, JPH3, CACNA2D2, SLC12A7, TNXB, OSTM1, SDK1) |
| Diabetes (Type 2) | basic (LRMDA)<br>bmi_9 (AMZ1, GNA12)<br>fathedu (SBNO2, HLA-DQB2)<br>momedu (NDUFAF6)<br>phys (HOXC4)<br>psyc (CRTAC1, CDKN1C, KCNJ11, CCND1, ANKK1, OPCML, |

| | |
|---|---|
| | HOXC4, WSCD2, NOS1, HHIPL1, ZNF280D, IGF1R, LMF1, CACNA1H, GSE1, KLHDC4, GLP2R, RAI1, SREBF1, MAPT-AS1, CSNK1G2, INHBB, LINC01101, NGEF, TCEA2, SCAP, CACNA2D2, IQCJ-SCHIP1, PDE6B, IDUA, JAKMIP1, EHMT2, CUTA, PACSIN1, ELFN1, HECW1, PRKAR2B, AGAP3)<br>smoke0 (CAT, LTBP3, KLHDC4, ZBTB38, DMGDH, POU5F1, ANK1, NDUFAF6) |
| Drug Use<br>Drug Dependence<br>Heroin Dependence<br>Opioid Dependence | bmi_9 (BRSK2, NCR1)<br>fdep (APOB)<br>mdep (GPRC5C)<br>psyc (CASZ1, MGMT, BET1L, LRP1, HHIPL1, SCHIP1, IQCJ-SCHIP1, SDK1, NOS3)<br>smoke0 (GABRB3) |
| Early Life Stress<br>Childhood Trauma | psyc (OSBPL5) |
| Eating Behavior | fdep (APOB)<br>psyc (NTM) |
| Educational Attainment<br>Self-Reported Educational Attainment | aces (TMEM232)<br>aces_threshold (TMEM232)<br>basic (LRMDA)<br>bmi_9 (BRSK2, IKZF4, FOXP4, TNXB)<br>bully9 (TYW3)<br>fathedu (ZNF512B, NRG2, HLA-DQB2, TNXB)<br>fdep (KLF6, APOB, TRPC3)<br>mdep (PRDM8, CD40)<br>momedu (HLA-DPA1)<br>nonv (NR4A2)<br>phys (PSMD9)<br>psyc (PRDM16, CRTAC1, SH3PXD2A, KNDC1, PC, NTM, OPCML, RPS27P25, BCL7A, ZNF280D, LMF1, CACNA1H, MAPT-AS1, FOXK2, CSNK1G2, MAP2K2, NR4A2, ALPI, NGEF, TCEA2, SUN2, CACNA2D2, SCHIP1, IQCJ-SCHIP1, RGS12, PRDM8, CAMK4, UNC5A, FLT4, COL11A2, GRM4, ELFN1, SDK1, HECW1, FLNC, PTPRN2, GPR20, RAI1, TRPC3, TNXB)<br>smoke0 (PRDM16, LTBP3, SHANK2, GABRB3, ZBTB38, ANK1) |
| Emotional Symptoms<br>Mood Instability<br>Mood Disorder | aces (TMEM232)<br>aces_threshold (TMEM232)<br>basic (LRMDA)<br>psyc (SSPN, MAPT-AS1) |
| Gestational Iron Deficiency | Basic (PMF125) |

| | |
|---|---|
| Insomnia | basic (LRMDA)<br>bmi_9 (GNA12)<br>fdep (ZFP57, PRDM1)<br>mdep (CD40)<br>psyc (PKD2L1, INPP5A, PC, NTM, LRP1, NOS1, IGF1R, KLHDC4, SLC38A10, ASPSCR1, CACNA2D2, CAMK4, ZFP57, PACSIN1, SDK1, PRKAR2B, CHD7)<br>smoke0 (KLHDC4, RFTN2) |
| Major Depressive Disorder | basic (LRMDA)<br>bmi_9 (TNXB, GNA12)<br>bully9 (TYW3, ZSCAN12P1)<br>fathedu (TNXB)<br>fdep (ZFP57)<br>mdep (PRDM8)<br>nonv (NR4A2)<br>psyc (CASZ1, SH3PXD2A, ANKK1, OPCML, SSPN, NOS1, PIPOX, MAPT-AS1, NR4A2, NGEF, TUBGCP6, PRDM8, ARHGEF37, ZFP57, TNXB, SDK1, BMPER, CHD7)<br>smoke0 (CAT, LTBP3, SHANK2, PIPOX, RFTN2, MYO1G) |
| Neonatal Lung Function<br>Childhood Lung Function | basic (C10orf1125) |
| Neuroticism | basic (LRMDA)<br>bmi_9 (TNXB, GNA12)<br>bully9 (TYW3)<br>fathedu (TNXB)<br>mdep (PM20D1, CD40)<br>momedu (HLA-DPA1)<br>nonv (NR4A2)<br>psyc (DIP2C, MGMT, ANKK1, OPCML, WSCD2, NOS1, SNHG14, RAI1, MAPT-AS1, NR4A2, SCAP, TNXB, SDK1) |
| Obsessive Compulsive Disorder | psyc (DRAXIN, RAI1, NGEF, IQCJ-SCHIP1, ANKK1, SREBF1) |
| Personality Disorder | psyc (SDK1)<br>mdep (CSNK1E24) |
| Psychotic Symptoms | psyc (DIP2C) |
| Schizophrenia | bmi_9 (BRSK2, TNXB, GNA12)<br>fathedu (NRG2, TNXB, HLA-DQB2)<br>fdep (KLF6, ZFP57, BRD2)<br>mdep (CD40)<br>nonv (NR4A2)<br>phys (NEAT1)<br>psyc (DRAXIN, MGMT, NTM, OPCML, LRP1, WSCD2, NOS1, |

| | |
|---|---|
| | ANKLE2, ADCY7, GSE1, JPH3, RAI1, NR4A2, GALNT5, NGEF, CACNA2D2, CD47, ZFP57, EHMT2, TNXB, TTYH3, SDK1, PTPRN2)<br>smoke0 (RFTN2, GTF2H4) |
| Smoking<br>Smoking Status<br>Smoking Initiation<br>Nicotine Use<br>Nicotine Dependence | basic (LRMDA)<br>bmi_9 (IKZF4, AMZ1, GNA12)<br>bully9 (SPTBN4)<br>fathedu (HLA-DQB2)<br>mdep (CD40, PRDM8)<br>phys (HOXC4)<br>psyc (TGFB2, NTM, OPCML, HOXC4, IGF1R, EHMT2, ELFN1, SDK1, PRDM16, CRTAC1, PC, SSPN, WSCD2, ZNF280D, SREBF1, PIPOX, MAPT-AS1, ASPSCR1, INHBB, LINC01101, NEU4, CD47, SCHIP1, IQCJ-SCHIP1, RBM46, ZNF316, SMARCD3, PTPRN2, SDCBP, CHD7, CASZ1, DIP2C, PRDM8, B3GALT4, NOS3)<br>smoke0 (ZBTB38, PRDM16, SHANK2, PIPOX, SEZ6) |
| Stress-related Disorders<br>Stress Reaction<br>Response to Trauma<br>PTSD | aces_threshold (GAREML/GAREM226)<br>basic (LRMDA)<br>mdep (CD40)<br>psyc (SDK1, MAPT-AS1, NOS1, CACNA2D2, ELFN1) |
| Substance Use<br>Substance Abuse<br>Substance Dependence | bully9 (TYW3)<br>mdep (CD40)<br>psyc (MGMT, NTM, WSCD2, IGF1R, ASPSCR1, IQCJ-SCHIP1, ELFN1, SDK1, PTPRN2, CHD7) |
| Suicide Behavior | psyc (OTUD7A, OPCML, BCL7A)<br>smoke0 (ENPEP) |
| Testosterone | bmi_9 (MFSD6L)<br>bully9 (VENTX)<br>fdep (BRD2)<br>psyc (TGFB2, SH3PXD2A, KNDC1, MIR202HG, HHIPL1, ZNF280D, RAI1, GREB1, ZNF316, SDCBP, CHD7)<br>smoke0 (NLGN2, RFTN2, VARS2) |
| Wellbeing Measurement | bmi_9 (TNXB)<br>bully9 (TYW3)<br>fathedu (TNXB, HLA-DQB2)<br>fdep (KLF6)<br>psyc (ANKK1, NOS1, TNXB, SDK1)<br>smoke0 (CMYA5) |

Table S4. Annotations of genes to DM in prior literature for genes that overlap with FFCWS ELA-associated DMRs across exposures.

| Conditions Associated with DM in Prior Literature | DMR Genes by FFCWS Exposure |
|---|---|
| Aggressive Behavior<br>Physically Aggressive Behavior | fathedu (*DRD4*[29])<br>momedu (*DRD4*[29]) |
| Alcohol Use Disorder<br>(Ventral Striatum) | aces (*TMEM232*[30])<br>aces_threshold (*TMEM232*[30]) |
| ADHD | momedu (*DRD4*[31])<br>fathedu (*DRD4*[31]) |
| Bullying | psyc (*HCG4*[32]) |
| Child Abuse<br>Physical Abuse<br>Physical or Emotional Abuse<br>Sexual Abuse | aces_threshold (*HOXA-AS3*[33])<br>bully9 (*VENTX*[34])<br>basic (TVP23A[35])<br>fathedu (*SBNO2*[33])<br>fdep (*HOXA-AS3*[33])<br>phys (*AKR7L*[36])<br>psyc (*NTM*[34], *SREBF1*[34], *PTPRN2*[34], *GALNT9*[36], *PRKCZ*[33], *PEX10*[33], *PTPRN2*[33], *SLC12A7*[33]) |
| Child Cognitive Deficits<br>Child Attention Deficits | fathedu (*DRD4*[37])<br>momedu (*DRD4*[37]) |
| Child Illness | bmi_9 (*TNXB*[36])<br>fathedu (*TNXB*[36])<br>psyc (*TNXB*[36]) |
| Childhood Obesity | bmi_9 (*TNXB*[38])<br>fathedu (*TNXB*[38])<br>mdep (*PRDM8*[38])<br>psyc (*TNXB*[38], *PRDM8*[38]) |
| Depression | psyc (*PTPRN2*[39], *TEKT4*[39])<br>smoke0 (*SAMD11*[39]) |
| Drug Addiction | fathedu (DRD4[40])<br>momedu (DRD4[40]) |
| Childhood Trauma<br>(Elderly individuals who experienced indentured childhood labor)<br>Childhood Neglect<br>(Physical Neglect)<br>Maltreatment | aces_threshold (*SKAP2*[41])<br>psyc (*CACNA1H*[41]) |
| Early Life Adversity<br>Childhood Stress Exposure<br>Early Life Stress in Animal Models | bmi_9 (*TNXB*[42], BRSK2[42], *ABAT*[42])<br>bully9 (*CRYZ*[42])<br>fathedu (*TNXB*[42]) |

| | |
|---|---|
| | fdep (*IFI44L*[42])<br>momedu (*HLA-DPB1*[42], *HLA-DPA1*[42])<br>phys (*WDR66*[43])<br>psyc (*PRDM16*[35,36,42], *TNXB*[42])<br>smoke0 (*CACNA2D4*[35], *GJD3*[35], *PRDM16*[35,36,42], *BHMT2*[42], *DMGDH*[42]) |
| Emotional Abuse<br>Emotional Neglect | bmi_9 (*ACY3*[36]) |
| Emotional Undereating in Childhood | aces (*CCDC169-SOHLH2*[44])<br>fathedu (*PF4*[44])<br>psyc (*PF4*[44]) |
| Exposure to Violence | nonv (*NR4A2*[45])<br>psyc (*NR4A2*[45])<br>smoke0 (*SHANK2*[45]) |
| Maternal Education | aces_threshold (*HOXA5*[46]) |
| Maternal Stressors<br>(Maternal Abuse)<br>(Maternal Sexual Abuse)<br>(Maternal Depression During Pregnancy)<br>(Maternal Physical Neglect)<br>(Maternal Neglect)<br>(Maternal Household Member Incarceration) | aces_threshold (*HOXA3*[47], *HOXA5*[46])<br>bmi_9 (*TNXB*[47], *MPRIP*[48])<br>fathedu (*TNXB*[47])<br>fdep (*HOXA3*[47], *TRPC3*[49], *ZFP57*[48])<br>momedu (*HLA-DPB1*[47])<br>phys (*BLCAP*[47]) |
| Neglect<br>Physical Neglect<br>Maltreatment | basic (*TVP23A*[35])<br>mdep (*PRDM8*[35])<br>psyc (*NTM*[35]) |
| Neighborhood Disadvantage | aces (*CCDC169*[33])<br>aces_threshold (*SKAP2*[33])<br>fdep (*TRPC3*[33])<br>momedu (*C17orf97*[33])<br>psyc (*ZNF280D*[33], *CSNK1G2*[33], *TRPC3*[33], *SPON2*[33], *CACNA2D2*[33]) |
| Nicotine Exposure During Pregnancy<br>(Fetal Tissue) | momedu (*C17orf97*)[21] |
| One-Adult Household | aces (*BOLL*[33])<br>psyc (*PDE6B*[33], *ELFN1*[33]) |
| Parental Stressors<br>(Parent Physical Illness)<br>(Parent Mental Illness)<br>(Maternal Psychopathology)<br>(Financial Stress) | aces_threshold (*C8orf31*[36])<br>bmi_9 (*C15orf26*[36], ABAT[33])<br>psyc (*PRDM16*[36], *DIP2C*[33], *CYP1B1*[33], *LRP1*[33], *TCF25*[33])<br>smoke0 (*PRDM16*[36]) |
| Paternal Stressors<br>(Father Depression)<br>(Expressed Anger)<br>(Parenting Stress) | aces_threshold (*HOXA3*[50])<br>fdep (*HOXA3*[49])<br>mdep (*CD40*[49])<br>momedu (*HLA-DPA1*[49]) |

| | |
|---|---|
| (Financial Stress during Preschool) | psyc (*OSBPL5*[49], *NEU4*[49]) |
| Post-Traumatic Stress Disorder (PTSD) | psyc (*AHRR*[50,51], *TOLLIP*[51])<br>smoke0 (*AHRR*[50,51]) |
| Prenatal Interventions and Infant Development (Low-Income Samples) | aces (*BOLL*)[52]<br>fathedu (*PF4*)[52]<br>momedu (*HLA-DPB1*)[52] |
| Prenatal Opioid Exposure<br>(Placental) | aces (*BOLL*)[53] |
| Primer for Stress Response<br>Stress-Resilience | aces (*SOHLH2*)[54]<br>bully9 (*CRYZ*[54]) |
| Psychotic Experiences<br>(Young Adulthood) | aces (*SOHLH2*[55]) |
| Schizophrenia<br>Sex-specific Schizophrenia | fathedu (*DRD4*[56,57])<br>momedu (*DRD4*[56,57]) |
| Smoking (Adult) | smoke0 (*MYO1G*[58–62], *AHRR*[58–69], *PIPOX*[64], *PRDM16*[64], *LTBP3*[62,64], *CNTNAP2*[62], *KLHDC4*[64]) |
| Smoking (Maternal Smoking in utero) | smoke0 (*ZBTB38*[70], *MYO1G*[11,71–74], *AHRR*[11,70,72–74]) |
| Social Anxiety Disorder | bmi_9 (*TNXB*[75])<br>fathedu (*TNXB*[75])<br>psyc (*TNXB*[75]) |
| Thyroid-stimulating Hormone<br>ft4 | aces (*CCDC169*[76])<br>bmi_9 (*UNC45A*[76])<br>bully9 (*VENTX*[76])<br>mdep (*UNC45A*[76]) |
| Time and Adversity Interaction | aces_threshold (*GAREML*[71])<br>phys (*GAREML*[71]) |

Table S5. Statistics of methylation-dependent regulatory associations by exposure, where beta represents the slope, p is the empirical p-value, and SE is the standard error.

| Exposure | beta | p | SE |
| --- | --- | --- | --- |
| aces | 0.001897 | 1.31E-02 | 0.000314 |
| aces_threshold | 0.001679 | 1.15E-03 | 0.000292 |
| basic | 0.00586 | 5.00E-05 | 0.000299 |
| bmi_9 | 0.00665 | 5.00E-05 | 0.000325 |
| bully9 | 0.020907 | 5.00E-05 | 0.000433 |
| fathedu | 0.00144 | 4.94E-02 | 0.000292 |
| fdep | -0.015223 | 1 | 0.000337 |
| mdep | 0.00407 | 5.00E-05 | 0.00027 |
| momedu | -0.006512 | 1 | 0.000351 |
| nonv | 0.000148 | 4.56E-01 | 0.00033 |
| phys | 0.014263 | 5.00E-05 | 0.000329 |
| psyc | 0.020551 | 5.00E-05 | 0.000586 |
| smoke0 | 0.000115 | 4.32E-01 | 0.000356 |

Table S6. Percent of DMRs lost, retained, and arising between Y9 and Y15.

| Exposure | %Y9 Persistent | %Y9 Only | %Y15 Retained | %Y15 Only |
|---|---|---|---|---|
| aces | 25 | 75 | 16.67 | 83.33 |
| aces_threshold | 66.67 | 33.33 | 100 | 0 |
| basic | 40 | 60 | 12.5 | 87.5 |
| bmi_9 | 30.43 | 69.57 | 50 | 50 |
| bully9 | 40 | 60 | 20 | 80 |
| fathedu | 53.85 | 46.15 | 36.84 | 63.16 |
| fdep | 20 | 80 | 22.22 | 77.78 |
| mdep | 50 | 50 | 38.46 | 61.54 |
| momedu | 30 | 70 | 12.5 | 87.5 |
| nonv | 33.33 | 66.67 | 50 | 50 |
| phys | 18.18 | 81.82 | 15.38 | 84.62 |
| psyc | 2.12 | 97.88 | 66.67 | 33.33 |
| smoke0 | 40 | 60 | 51.43 | 48.57 |

### Supplemental Figures

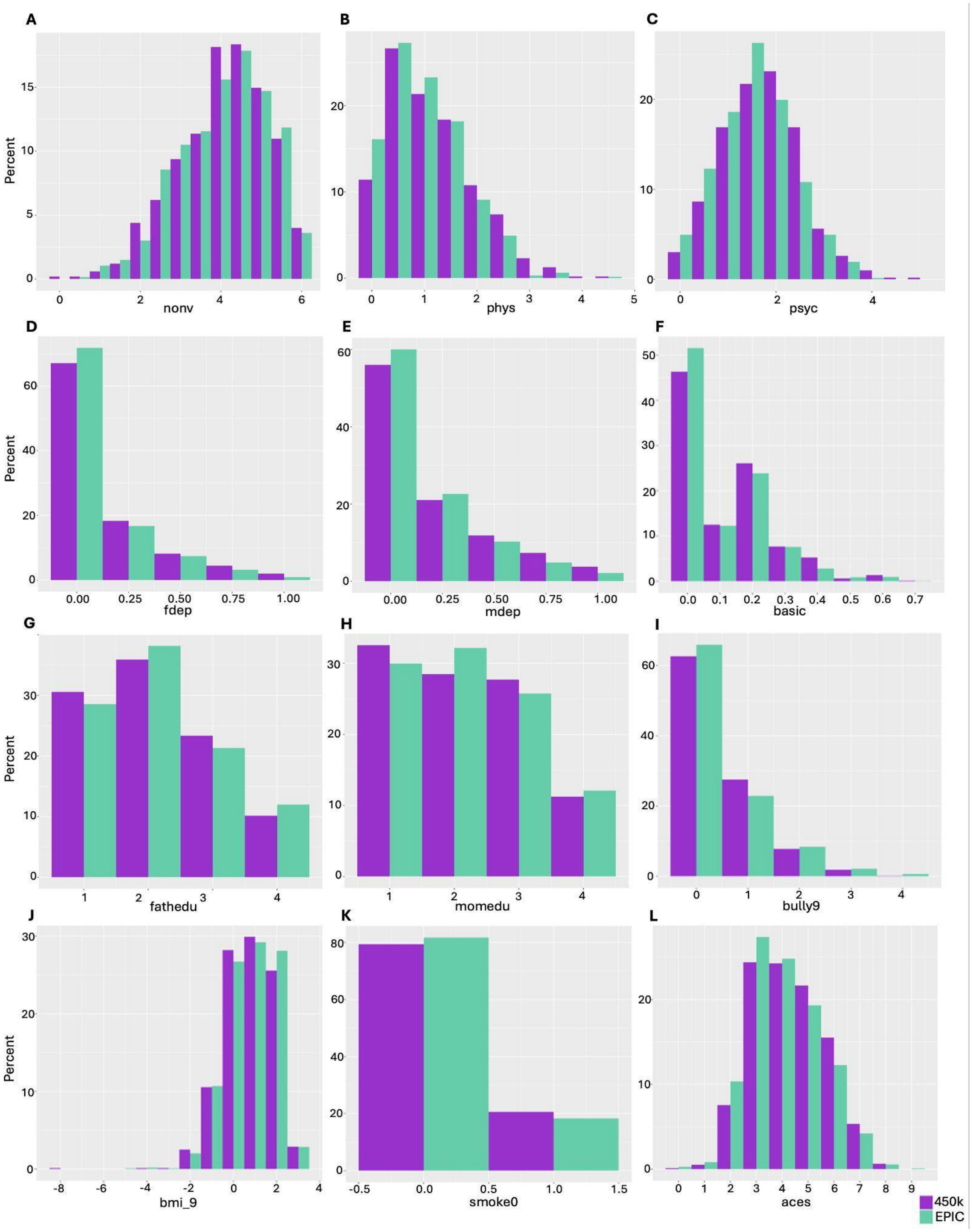


**Figure S1: Exposure distribution scores for the 450K and EPIC arrays.** Phenotype distribution for the 450K array in purple and EPIC array in green for (A) nonviolent discipline; (B) physical assault; (C) psychological aggression; (D) paternal depression; (E) maternal depression; (F) material hardship; (G) father education level; (H) maternal education level; (I) peer bullying; (J) BMI at age 9; (K) maternal smoking during pregnancy; (L) cumulative ACEs.

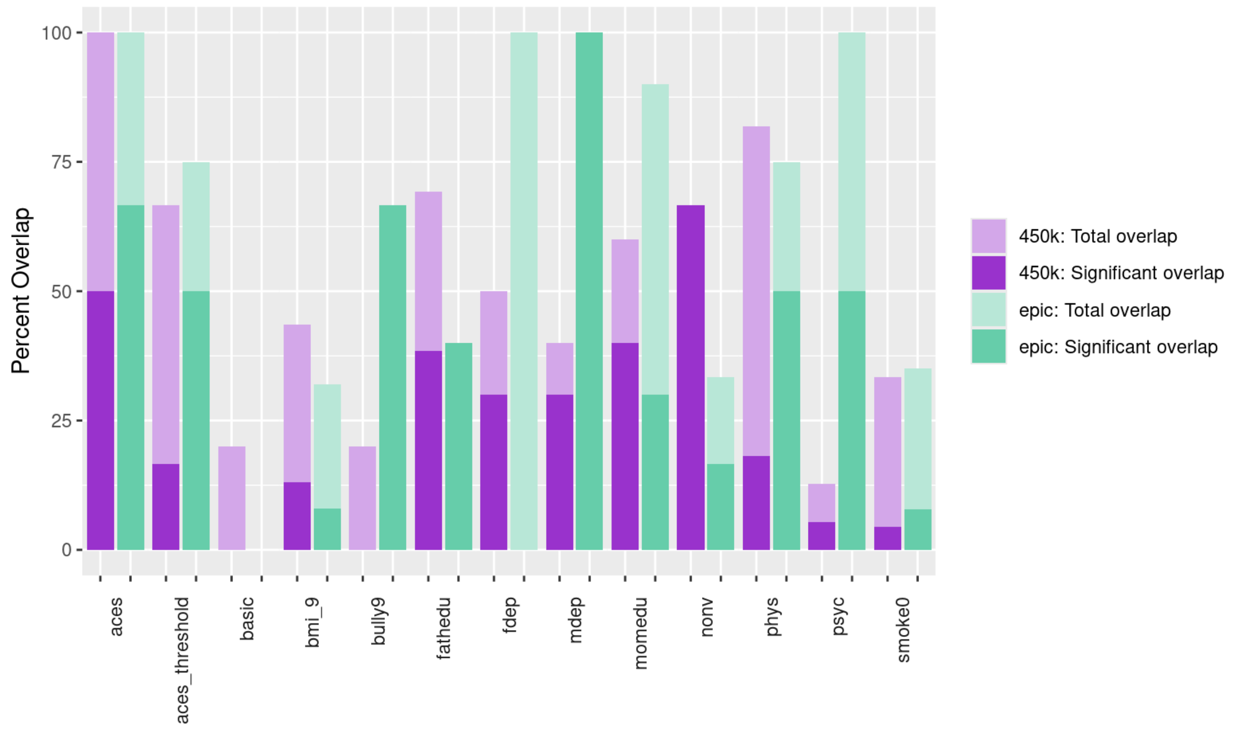


**Figure S2**: **Percent of DMRs that overlap with another DMR across 450K and Epic arrays.** Darker purple/green indicates an overlap with significant DMRs (pval < 0.001) for other phenotypes and lighter purple/green indicates an overlap with a DMR that has a region corrected p-value < 0.05.

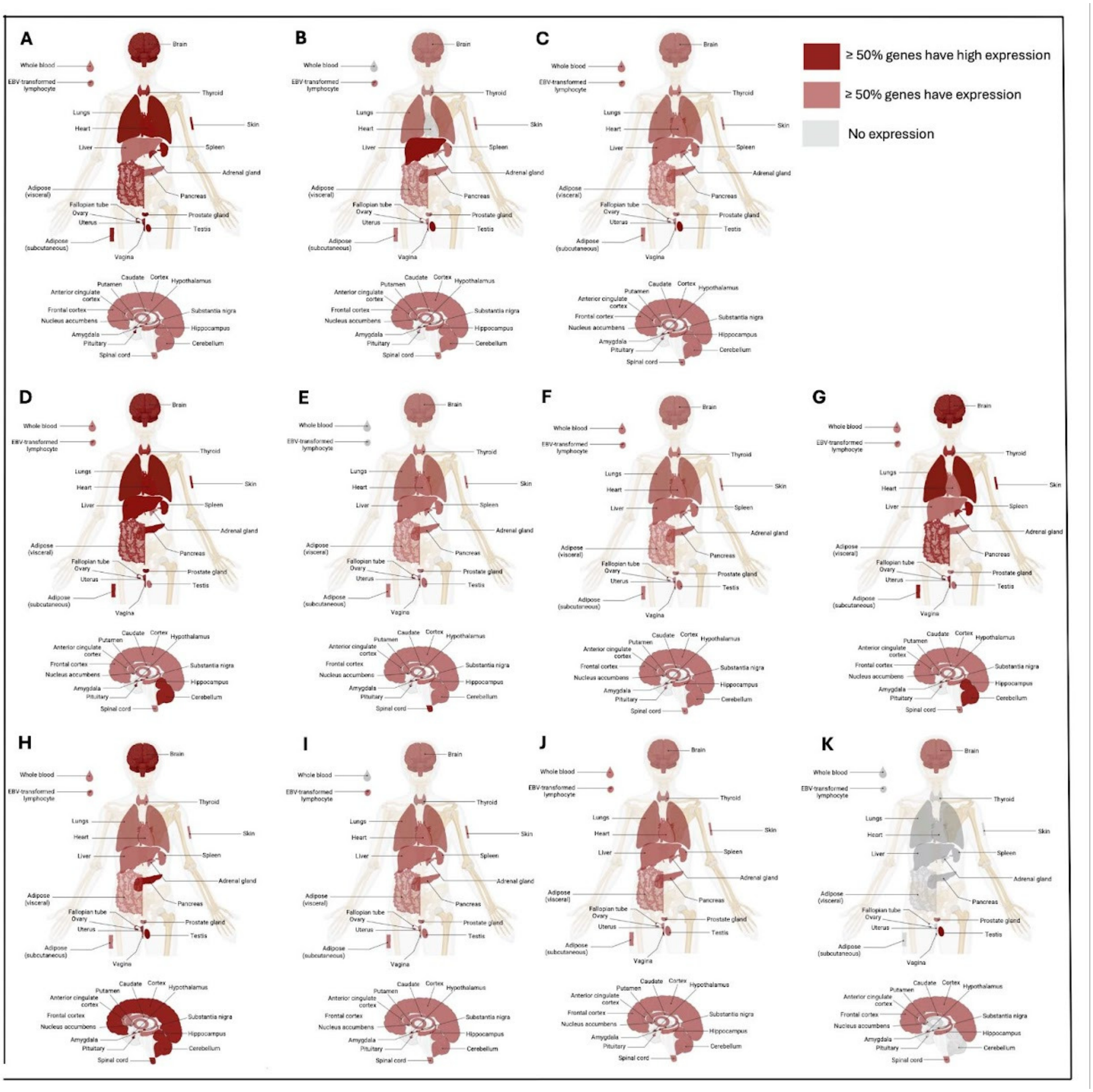


**Figure S3: Tissue-specific expression of genes overlapping 450K array DMRs for additional exposures.**

GTEx expression of genes overlapping significant DMRs for (A) *nonv*: nonviolent discipline; (B) *phys*: physical assault; (C) *psyc*: psychological aggression; (D) *mdep*: maternal depression; (E) *fdep*: paternal depression; (F) *momedu*: mother education level; (G) *fathedu*: father education level; (H) *bmi_9*: BMI at age 9; (I) *smoke0*: maternal smoking during pregnancy; (J) *basic*: material hardship; (K) *aces*: cumulative ACEs. Light red indicates that at least 50% of genes are expressed in a given tissue; dark red indicates at least 50% of genes have high expression (median TPM ≥ 10) in a given tissue.

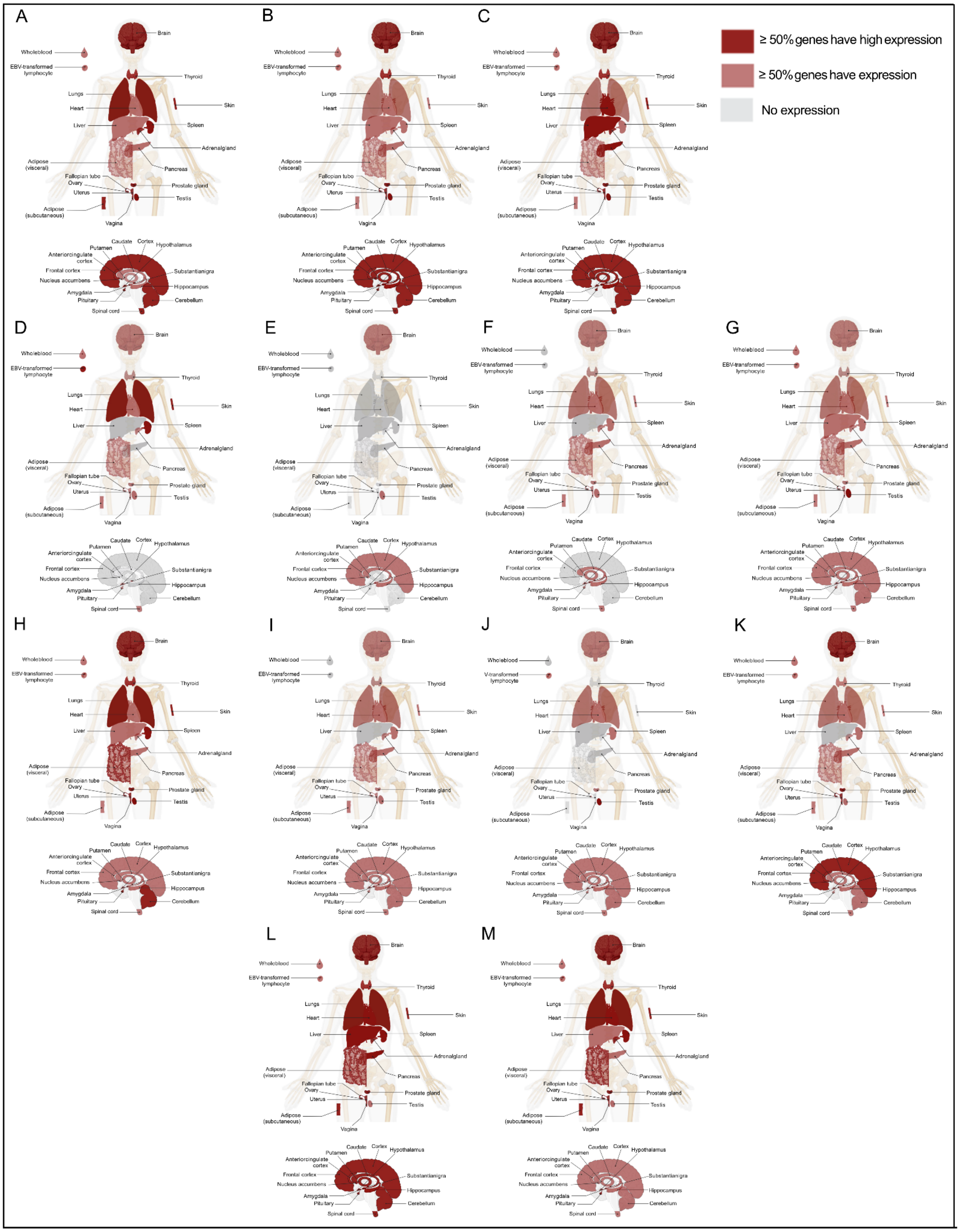


**Figure S4: Tissue-specific expression of genes overlapping EPIC array DMRs for additional exposures**. GTEx expression of genes overlapping significant DMRs for (A) nonviolent discipline; (B) physical assault; (C) psychological aggression; (D) maternal depression; (E) paternal depression; (F) mother education level; (G) father education level; (H) BMI at age 9; (I) maternal smoking during pregnancy; (J) material hardship; (K) peer bullying; (L) cumulative ACEs; (M) thresholded cumulative ACEs. Light red indicates that at least 50% of genes are expressed in a given tissue; dark red indicates at least 50% of genes have high expression (median TPM ≥ 10) in a given tissue. * indicates that expression is greater than expected by chance (Wilcoxon ranked-sum test) relative to the full gene-set included in GTEx ($q < 0.05$).

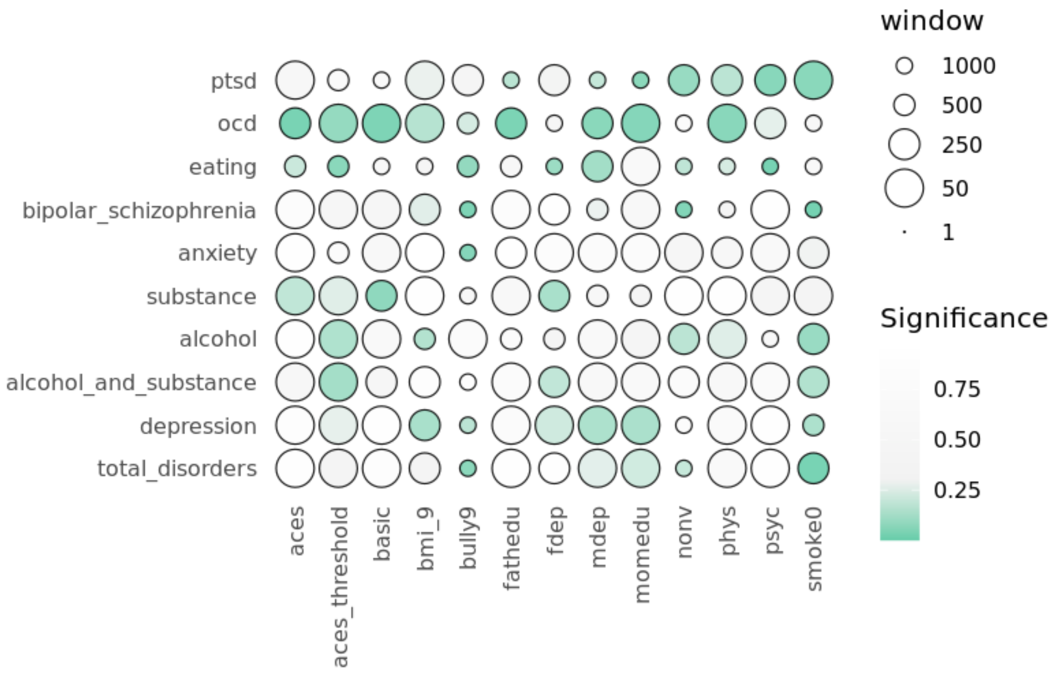


**Figure S5: Association of differential methylation with nearby SNPs and associated disorders (EPIC array).** Window for exposure-associated DM near a disease-associated SNP (within 50/250/500/1000bp) is represented as circle size inversely proportional to window size, such that closer DM-SNP pairs occupy larger circles. Significance was determined using empirical p-values, with $p_{emp}$<0.3 shown as a pink color gradient.

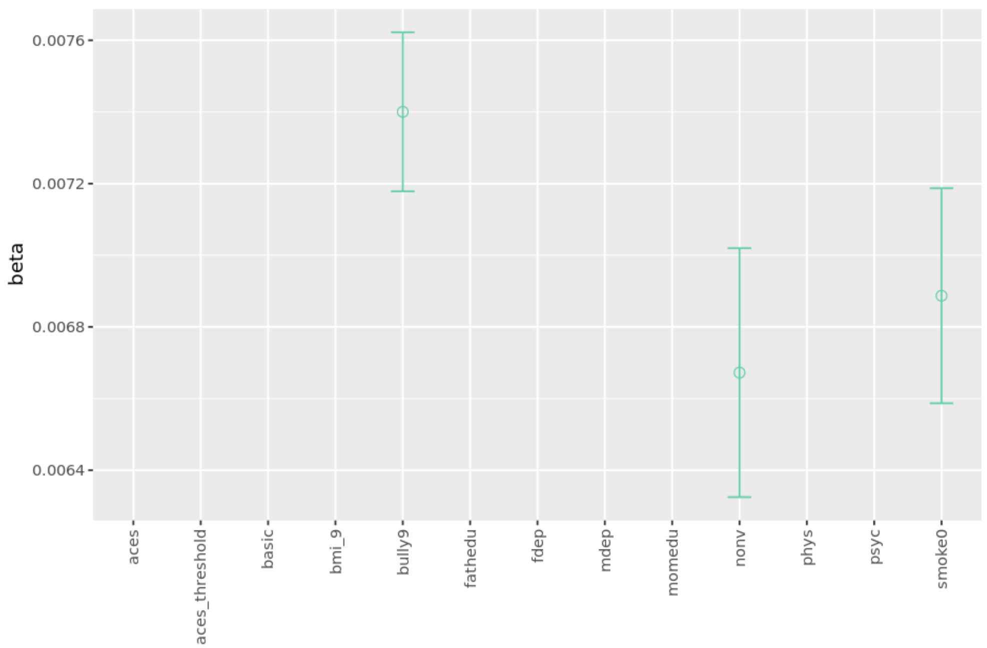


**Figure S6: Relationship between differential methylation and methylation-dependent gene regulatory activity for the EPIC array.** Positive Beta coefficients for each regression between DM significance and methylation-dependent regulatory activity slopes ($\beta$, Equation 3) for each exposure. No beta slopes met statistical significance (*p-value < 0.05*).

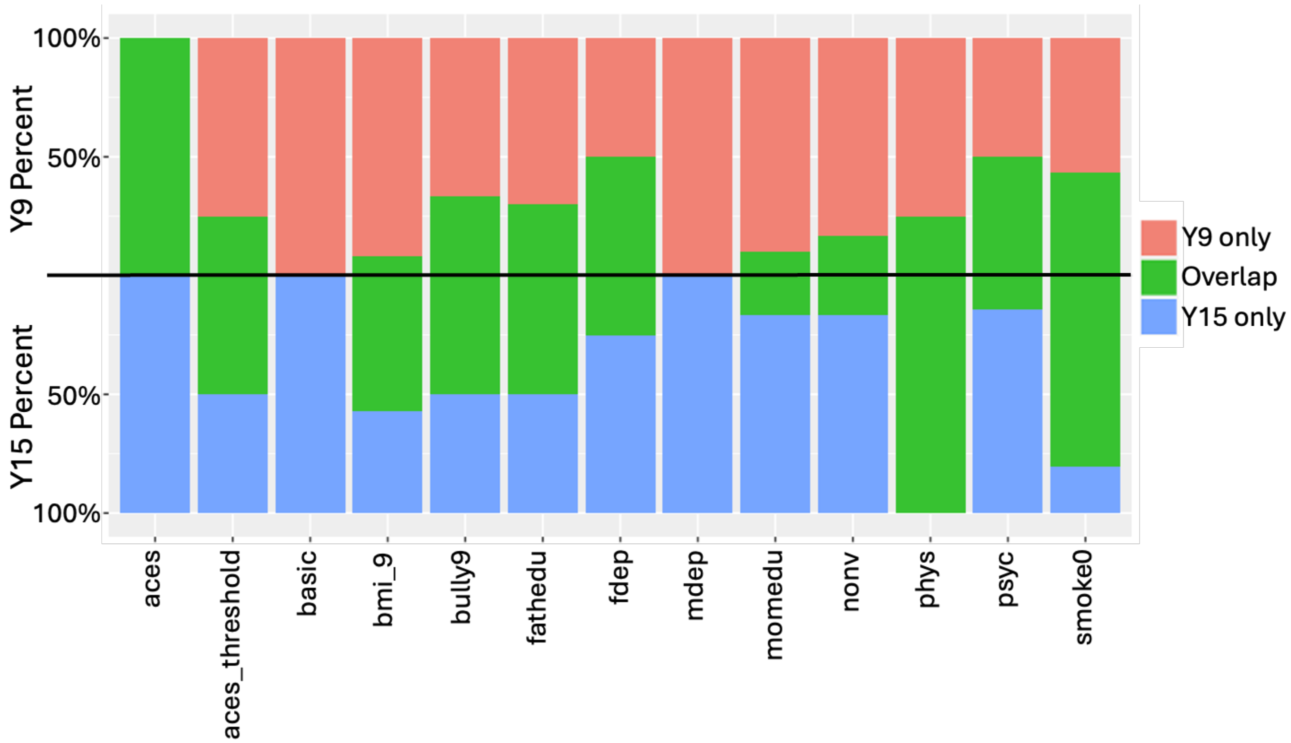


**Figure S7: Percent of DMRs lost, retained, or added between ages 9 and 15 for the EPIC array.**
Values on top reflect percent of Y9 DMRs specific to Y9 (red) or overlapping with Y15 (green). Values on bottom reflect percent of Y15 DMRs specific to Y15 (blue) or overlapping with Y9 (green). Each exposure is represented as a separate bar.

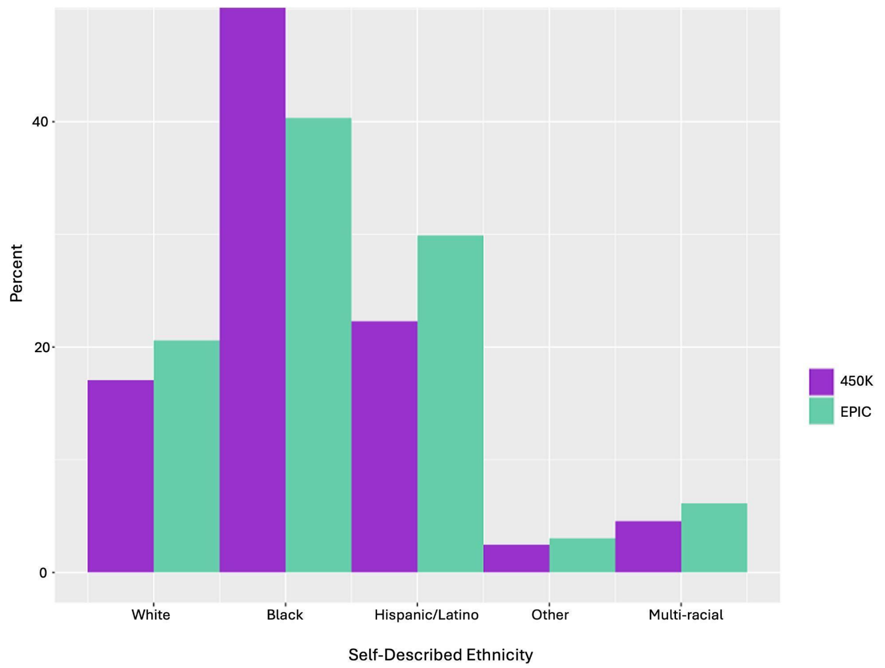


**Figure S8: Percent of self-described ethnicity across participants on 450K and EPIC arrays.** Histogram of self-described ethnicity across 767 participants on the 450K array (purple) and 1093 participants on the EPIC array (green). White self-reported as white only, non-hispanic; Black as Black/African American only, non-hispanic; Other as Other-only, non-hispanic, Multi-racial as Multi-racial, non-hispanic; or Hispanic/Latino.
